# AUTS2, a causative gene for microcephaly, regulates division of intermediate progenitor cells and production of upper-layer neurons

**DOI:** 10.1101/2024.04.15.589462

**Authors:** Kazumi Shimaoka, Kei Hori, Satoshi Miyashita, Yukiko U. Inoue, Asami Sakamoto, Ikuko Hasegawa, Kunihiko Yamashiro, Saki F. Egusa, Shoji Tatsumoto, Yasuhiro Go, Manabu Abe, Kenji Sakimura, Takayoshi Inoue, Takuya Imamura, Mikio Hoshino

## Abstract

*AUTS2* mutations often exhibit neurodevelopmental disorders and microcephaly. However, how AUTS2 regulates neuron production and affects brain size remains unclear. Here, we show that AUTS2 cooperates with the Polycomb complex PRC2 to regulate gene expression and cortical neurogenesis. *Auts2* mutant mice exhibit reduced division of intermediate progenitor cells (IPCs), leading to decreased neurons and thickness in the upper-layer cortex. Expression of *Robo1* is increased in the mutants, which in turn suppresses IPC division. Transcriptome and chromatin profiling experiments show that, in IPCs, AUTS2 primarily represses transcription of genes, including *Robo1*. Promoter region of AUTS2 target genes is enriched with H3K27me3, a repressive histone modification, but its levels are reduced in *Auts2* mutants. AUTS2 interacts with PRC2, and together, they promote IPC division. These suggest that AUTS2 collaborates with PRC2 to repress gene transcription through H3K27 trimethylation, promoting neuron production. This sheds light on AUTS2 pathophysiological mechanisms in neurogenesis and microcephaly.

## Introduction

*AUTS2* has been identified as a possible autism spectrum disorders (ASD) risk gene in a pair of monozygotic twins^1^. Structural variants and single nucleotide polymorphisms (SNPs) in the *AUTS2* locus have since been reported as associated with various psychiatric disorders^2,3^. Patients with the AUTS2 syndrome carry heterozygous mutations and exhibit a high frequency of intellectual disability (ID) (98%), as well as attention deficit hyperactivity disorder (ADHD) and ASD^4–6^. In addition, several common pathological features have been reported frequently in the AUTS2 syndrome, particularly that 65% of the patients present with microcephaly^4,6,7^. Recently, magnetic resonance imaging (MRI) analysis has reported cortical volume loss, cerebellar hypoplasia, and corpus callosum (CC) hypoplasia in patients with AUTS2 syndrome^8,9^. *Auts2*-deficient mice show behavioral abnormalities that mimic some of the human symptoms, such as impaired memory and learning and social behavior^10–13^. Notably, the brains of the mice have also been reported to exhibit cerebellar hypoplasia^14^, which may recapitulate the cerebellar hypoplasia seen in patients with AUTS2 syndrome. However, hypoplasia of the cerebral cortex corresponding to microcephaly, which is seen in many patients, has not been reported in mouse models. In zebrafish, *auts2a* knockdown (KD) by morpholinos has been shown to result in small brain^7,15^. Cerebral organoids derived from an AUTS2 syndrome patient exhibited reduced growth^8^. Although microcephaly may be caused by a decrease in the number of neurons, the mechanism by which microcephaly is caused as a result of impaired AUTS2 function remains unclear. Therefore, it is necessary to recapitulate the phenotype in a mouse model and analyze it to elucidate the molecular mechanism *in vivo*.

In the mammalian cerebral cortex, neurons are generated directly from progenitor cells called radial glial cells (RGCs) on the ventricular surface during early neurogenesis (direct neurogenesis)^16^. These neurons differentiate primarily into deep-layer neurons^17^. As development proceeds, RGCs generate secondary progenitors called IPCs, which divide once to several times in the subventricular zone (SVZ), eventually producing 2-12 neurons (indirect neurogenesis)^18–21^. Neurons derived from IPCs differentiate mainly into upper-layer neurons^22,23^. Mammals have newly acquired IPCs during evolution, which has contributed to the expansion of the mammalian cerebral cortex^24^. While the mechanisms of RGC proliferation and differentiation into neurons have been well investigated^25,26^, the regulatory mechanisms of IPC proliferation and differentiation remain largely unexplored.

Previous studies have reported that AUTS2 is localized and functions in cell nuclei and cytoplasm^13,27^. Cytoplasmic AUTS2 regulates the actin cytoskeleton through the RAC1 signaling pathway, which plays a pivotal role for neuronal migration and neurite elongation^27^. Nuclear AUTS2 limits the number of synapses and regulates the E/I balance in the brain^10^. It has been reported that nuclear AUTS2 acts as a transcriptional activator or repressor, depending on the cell type^9,12,13,28^. Gao et al. have shown the transcriptional activation machinery that AUTS2 interacts with non-canonical polycomb repressive complex 1 (PRC1.3/5) and recruits histone acetyltransferase P300 to acetylate at H3K27^13^. However, the molecular mechanism of how AUTS2 is involved in transcriptional repression is not well understood.

In this study, we have successfully generated mouse models of AUTS2 syndrome that recapitulate microcephaly. In addition, we discovered a new mechanism that regulates IPC proliferation and differentiation. We also found that AUTS2 interacts with PRC2 in the nucleus of IPCs, which affects chromatin modifications and represses gene transcription. This study not only leads to a better understanding of the pathogenesis of microcephaly in AUTS2 syndrome and the molecular function of AUTS2, but also sheds light on the evolution of the mammalian brain.

## Results

### Loss of *Auts2* leads to the reduction of upper-layer neurons in the cerebral cortex

We generated forebrain-specific *Auts2* conditional knockout (*Auts2* cKO) homozygous and heterozygous mice (*Emx1^Cre/+^; Auts2^flox/flox^*, *Emx1^Cre/+^; Auts2^flox/+^*) by crossing *Auts2-floxed* mice with *Emx1^Cre^*mice to investigate the role of AUTS2 in cerebral corticogenesis^10^. We previously confirmed that the full-length AUTS2 (FL-AUTS2) and C-terminal AUTS2 short isoform variant 1 (S-AUTS2-var.1) were successfully eliminated in the forebrains of *Auts2* cKO homozygotes^10^. In the Nissl staining of postnatal day (P) 30 brain sections, we did not observe apparent morphological differences between the control and *Auts2* cKO homozygous cortices at the macroscopic level (Figure 1A). However, upon closer observation, the cerebral cortex of cKO homozygotes appeared thinner than that of the controls (Figure 1B). Next, we performed immunostaining with layer-specific neuronal markers in coronal sections of the P15 control, heterozygous, and homozygous cortices at three points along the rostrocaudal axis. Notably, the number of upper-layer neurons (CUX1^+^ cells) was lower in homozygotes and heterozygotes than in controls (Figures 1C and D). This tendency was more severe in rostral sections than in caudal sections. In contrast, no significant differences were observed in the number of deep-layer neurons (CTIP2^+^ cells) among genotypes (Figure 1D). The upper-layer cortex was thinner in heterozygotes and homozygotes than in controls at the rostral point, whereas no significant differences were observed at the central or caudal points (Figures 1C and E). We also observed that the thickness of the rostral deep-layer cortex was slightly decreased in homozygotes compared with that in controls (Figure 1E).

**Figure 1:**
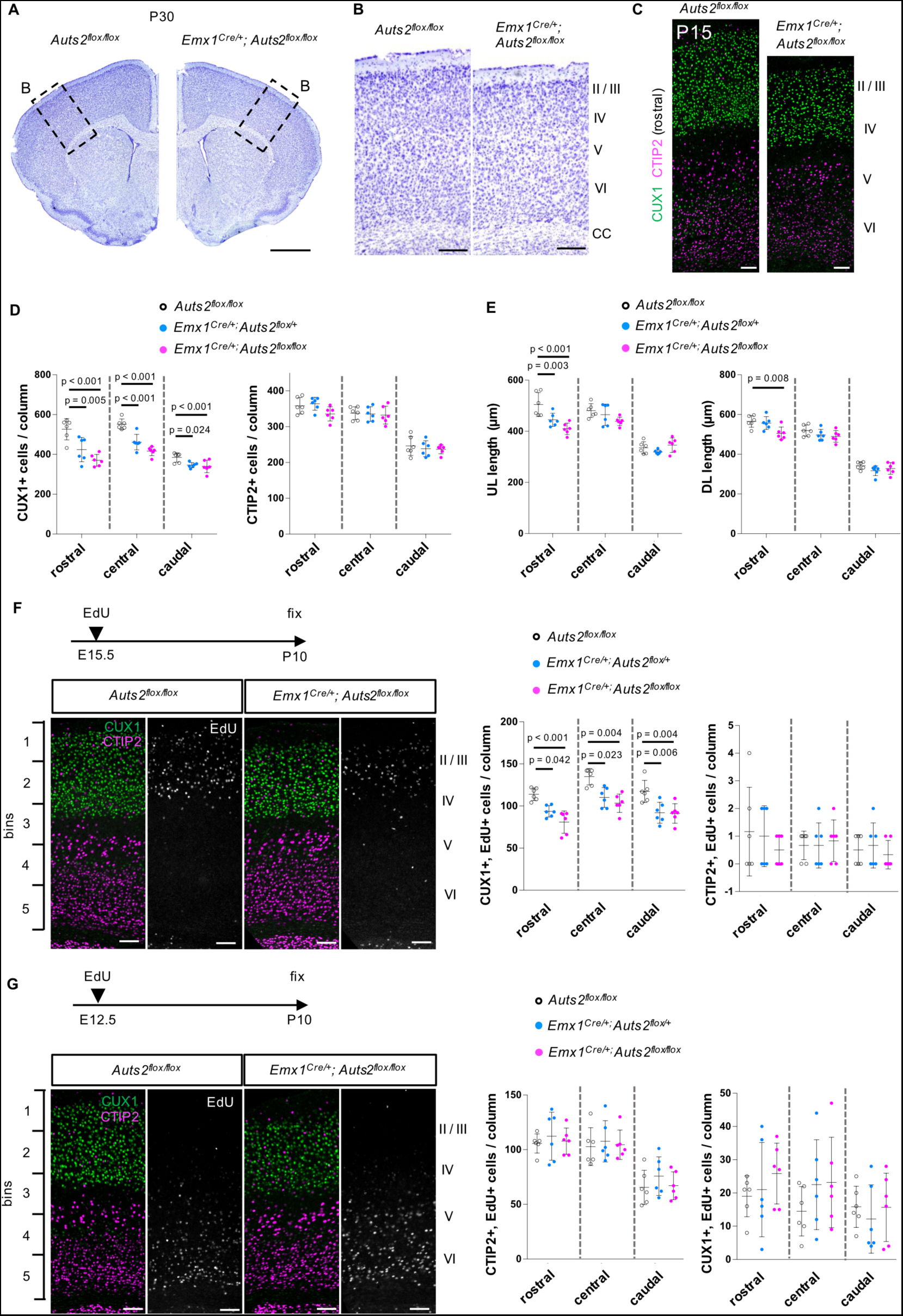
AUTS2 regulates the production of upper-layer neurons. (A) Nissl-stained sections of P30 brain hemispheres of indicated genotypes. (B) High magnification pictures of rectangle regions in (A), showing the somatosensory cortex. (C) Representative images of immunostaining for CUX1 (green) and CTIP2 (magenta) on the coronal sections at the rostral point of the cerebral cortex in *Auts2^flox/flox^* and *Emx1^Cre/+^; Auts2^flox/flox^*at P15. (D) Quantification of the number of CUX1^+^ (left) or CTIP2^+^ (right) cells in *Auts2^flox/flox^*, *Emx1^Cre/+^; Auts2^flox/+^* and *Emx1^Cre/+^; Auts2^flox/flox^* mice at P15 within a 300 µm wide field. (E) Quantification of the thickness of upper- or deep-layer in *Auts2^flox/flox^*, *Emx1^Cre/+^; Auts2^flox/+^* and *Emx1^Cre/+^; Auts2^flox/flox^* mice at P15. (F) EdU was administered intraperitoneally once to E15.5 pregnant mice. Representative images of triple-staining for CUX1 (green), CTIP2 (magenta), and EdU (white) in the cerebral cortices from *Auts2^flox/flox^* and *Emx1^Cre/+^; Auts2^flox/flox^* mice at P10. The graphs show the number of CUX1^+^/BrdU^+^ (left) and CTIP2^+^/EdU^+^-double positive cells (right) in *Auts2^flox/flox^*, *Emx1^Cre/+^; Auts2^flox/+^* and *Emx1^Cre/+^; Auts2^flox/flox^* mice within 300 µm wide field, respectively. (G) EdU was administered intraperitoneally once to E12.5 pregnant mice. Representative images of triple-staining for CUX1 (green), CTIP2 (magenta), and EdU (white) in the cerebral cortices from *Auts2^flox/flox^* and *Emx1^Cre/+^; Auts2^flox/flox^* mice at P10. The graphs show the number of CTIP2^+^/BrdU^+^ (left) and CUX1^+^/EdU^+^-double positive cells (right) within a 300 µm wide field, respectively. Data are presented as the mean ± SD (N = three mice, six sections); One-way ANOVA with Dunnett’s post-hoc test or Kruskal–Wallis test. Scale bars, 1 mm (A), 200 µm (B), and 100 µm (C, F, and G).

We did not find an increased number of cells with cleaved caspase-3, an apoptotic cell death marker, in the cerebral cortex of *Auts2* cKO mice at the embryonic and postnatal stages (Figure S1A), suggesting that the reduced size of the cerebral cortex or reduced number of upper-layer neurons in *Auts2* cKO mice was not likely due to cell death. Next, we performed pulse-labeling with 5-ethynyl-2’-deoxyuridine (EdU) on embryos at embryonic days (E) 15.5 and E12.5, when the upper- and deep-layer neurons were preferentially generated, respectively^17^. At P10, the number of upper-layer neurons (CUX1^+^ cells) labeled with EdU at E15.5 was significantly reduced throughout the cortices of *Auts2* cKO homozygotes and heterozygotes compared with *Auts2^flox/flox^* controls at P10 (Figure 1F). This suggests that upper-layer neurons derived from neural progenitors at E15.5 were reduced at P10 in *Auts2* cKO homozygotes and heterozygotes. In contrast, the number of CTIP2^+^ cells labeled with EdU at E12.5 was not significantly different among the genotypes (Figure 1G). As expected, we found fewer E15.5-labeled CTIP2^+^ neurons and E12.5-labeled CUX1^+^ neurons, with no significant differences among the genotypes (Figures 1F and G). A reduction in E15.5-labeled upper-layer neurons (CUX1^+^ cells) was observed as early as E18.5 in homozygotes and heterozygotes for the conventional *Auts2* allele (*Auts2^del8^*)^27^ (Figure S1B). These observations suggest that AUTS2 is required for upper-layer neuron production from neural progenitors.

We previously observed that most E14.5-labeled neurons did not reach the pial side by E18.5 in *Auts2* KO mice, whereas those in wild-type (WT) mice reached this level, suggesting that AUTS2 is involved in neuronal migration^27^. However, we found that the distribution of E15.5-labeled and E12.5-labeled cells did not differ between controls and homozygotes or heterozygotes at P10 (Figure S1C). This suggests that *Auts2*-deficient neurons migrated slowly but eventually reached their final targets. A few CUX1^+^ neurons were consistently observed in the deep layers (V and VI) of *Auts2* cKO homozygotes at P15 (Figure 1C).

Bedogni et al. reported that in *in situ* hybridization experiments, *Auts2* transcripts were observed in radial glial cells (RGCs) and IPCs^29^. By analyzing previous single-cell RNA-sequencing (scRNA-seq) data from the cerebral cortices at E13.5 and E15.5^30^, we found that *Auts2* transcripts were expressed in postmitotic neurons, RGCs, and IPCs (Figures 2A-F). These findings suggest that AUTS2 is expressed in neural progenitors (RGCs and IPCs) in the developing cerebral cortex; however, it might be below the detection level in these cells in a previous immunohistochemical experiment^27^.

**Figure 2:**
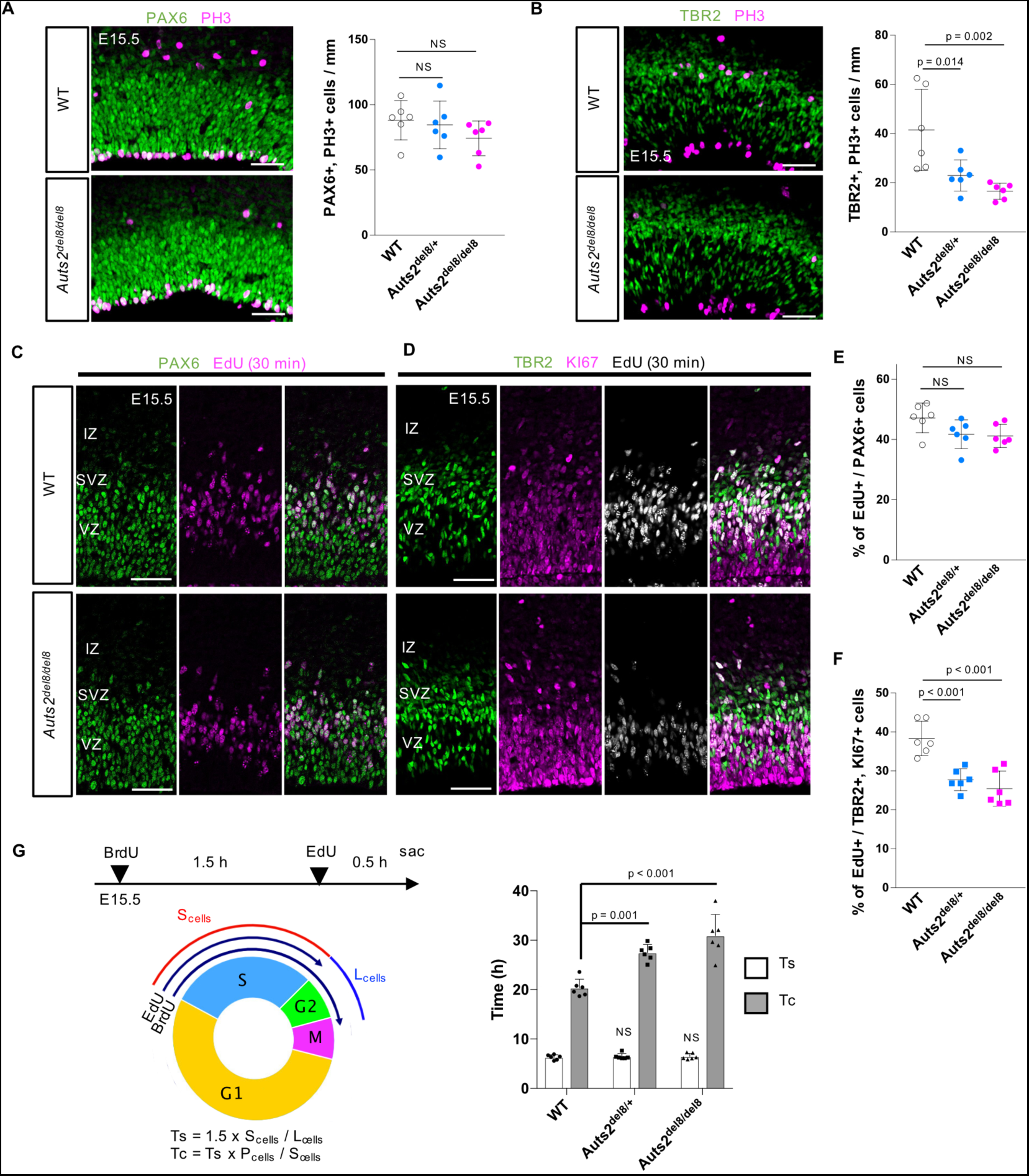
Loss of *Auts2* leads to the reduction of intermediate progenitor mitoses. (A) Immunostaining for PAX6 (green) and PH3 (magenta) in E15.5 WT and *Auts2^del8/del8^* cerebral cortices. The graph shows the number of PAX6^+^ and PH3^+^ cells on the ventricular surface in WT, *Auts2^del8/+^*, and *Auts2^del8/del8^*mice. (B) Immunostaining for TBR2 (green) and PH3 (magenta) in E15.5 WT and *Auts2^del8/del8^* cerebral cortices. The graph shows the number of TBR2^+^ and PH3^+^ cells in WT, *Auts2^del8/+^*, and *Auts2^del8/del8^*mice. (C–F) EdU was administered intraperitoneally to pregnant mice 30 min before sacrifice at E15.5. (C) Representative images of staining for PAX6 (green) and EdU (magenta) in WT and *Auts2^del8/del8^* cortical sections. (D) Representative images of staining for TBR2 (green), KI67 (magenta), and EdU (white) in WT and *Auts2^del8/del8^* cortical sections. (E, F) The graphs show the percentage of EdU^+^ cells among PAX6^+^ cells (E) or TBR2^+^ KI67^+^ cells (F) in WT, *Auts2^del8/+^*and *Auts2^del8/del8^* mice. VZ, ventricular zone; SVZ, subeventricular zone; IZ, intermediate zone. (G) The cell-cycle length estimate in TBR2^+^ cells using the BrdU and EdU double-labeling method at E15.5. BrdU and EdU are administered to pregnant mice at 2 h and 30 min before sacrifice. Ts, S-phase length; Tc, cell cycle length; L_cells_, BrdU^+^, and EdU-negative cells; S_cells_, EdU^+^ cells; P_cells_, TBR2^+^ cells. The number of cells was quantified at the rostral point. Data are presented as the mean ± SD (N = three mice, six sections). NS, not significant, One-way ANOVA with Dunnett’s post-hoc test. Scale bars, 50 µm.

### AUTS2 is involved in the proliferation of TBR2^+^ intermediate progenitor cells

Immunostaining of the E15.5 cerebral cortices with PAX6 antibody, a marker for RGCs, showed that the number of RGCs was not affected in *Auts2^del8/+^*or *Auts2^del8/del8^* mice (Figure S3A). Englund et al. previously showed that TBR2 (gene: *Eomes*)-positive cells in the developing cerebral cortex are largely IPCs^31^; however, they include a small population of immature neurons that have just emerged from IPCs. Immunostaining showed that approximately 85% of TBR2^+^ cells were KI67^+^ in the WT cerebral cortex at E15.5 (Figure S3B), indicating that most TBR2^+^ cells were IPCs. The number of TBR2^+^ cells or TBR2/KI67-double positive cells was not affected in *Auts2* heterozygotes (*Auts2^del8/+^*) or homozygotes (*Auts2^del8/del8^*) (Figure S3B). These findings indicated that the number of neural progenitors (RGCs and IPCs) did not change in the mutants, leading us to investigate the proliferation rate of neural progenitors.

Co-immunostaining for PAX6 and phosphorylated histone H3 (PH3), a mitotic (M)-phase cell marker, to E15.5 cortices showed that the number of dividing RGCs at the ventricular surface did not differ among the genotypes (Figure 2A). In contrast, the number of mitotic IPCs (TBR2^+^ and PH3^+^ cells) was markedly reduced in *Auts2* heterozygotes (*Auts2^del8/+^*) and homozygotes (*Auts2^del8/del8^*) (Figure 2B). Moreover, short-pulse labeling with EdU revealed that the proportion of IPCs (TBR2^+^ and KI67^+^ cells) in the synthesis phase (S-phase) was significantly decreased in *Auts2* heterozygotes (*Auts2^del8/+^*) or homozygotes (*Auts2^del8/del8^*) at E15.5, whereas that of RGCs (PAX6^+^ cells) was comparable between the WT and *Auts2* mutants (Figures 2C– F). At an earlier embryonic stage (E12.5), we observed no significant differences in the number of M-phase (PH3^+^) and S-phase (acute EdU^+^) cells in either the PAX6^+^ or TBR2^+^ population between the WT and *Auts2* mutants (Figure S3C–J). Given the observation that the number of IPCs did not differ among genotypes (Figure S3B) and that IPCs in the M-phase and S-phase were both reduced in *Auts2* heterozygotes and homozygotes at E15.5, we assumed that the cell cycle length of IPCs might be prolonged in *Auts2* mutants at this stage.

We then investigated the cell cycle of TBR2^+^ cells at E15.5, using the dual-labeling method with 5-bromo-2′-deoxyuridine (BrdU) and EdU^32^ and found that the length of the overall cell cycle (Tc) of TBR2^+^ cells was significantly longer in *Auts2* heterozygotes (*Auts2^del8/+^*) and homozygotes (*Auts2^del8/del8^*) than in WT mice. However, the S-phase length (Ts) was comparable among the genotypes (Figure 2G). Since a large population of TBR2^+^ cells were IPCs (Figure S3B), and since the number of IPCs (TBR2^+^ and KI67^+^ cells) was not affected in *Auts2* mutants (Figure 3B), it was suggested that the cell cycle length of IPCs was longer in *Auts2* mutants than that in WT mice. Consequently, these findings indicate that IPCs in the mutants divide less frequently than those in WT mice and that AUTS2 may be involved in the proliferation of IPCs and the subsequent production of upper-layer neurons.

**Figure 3:**
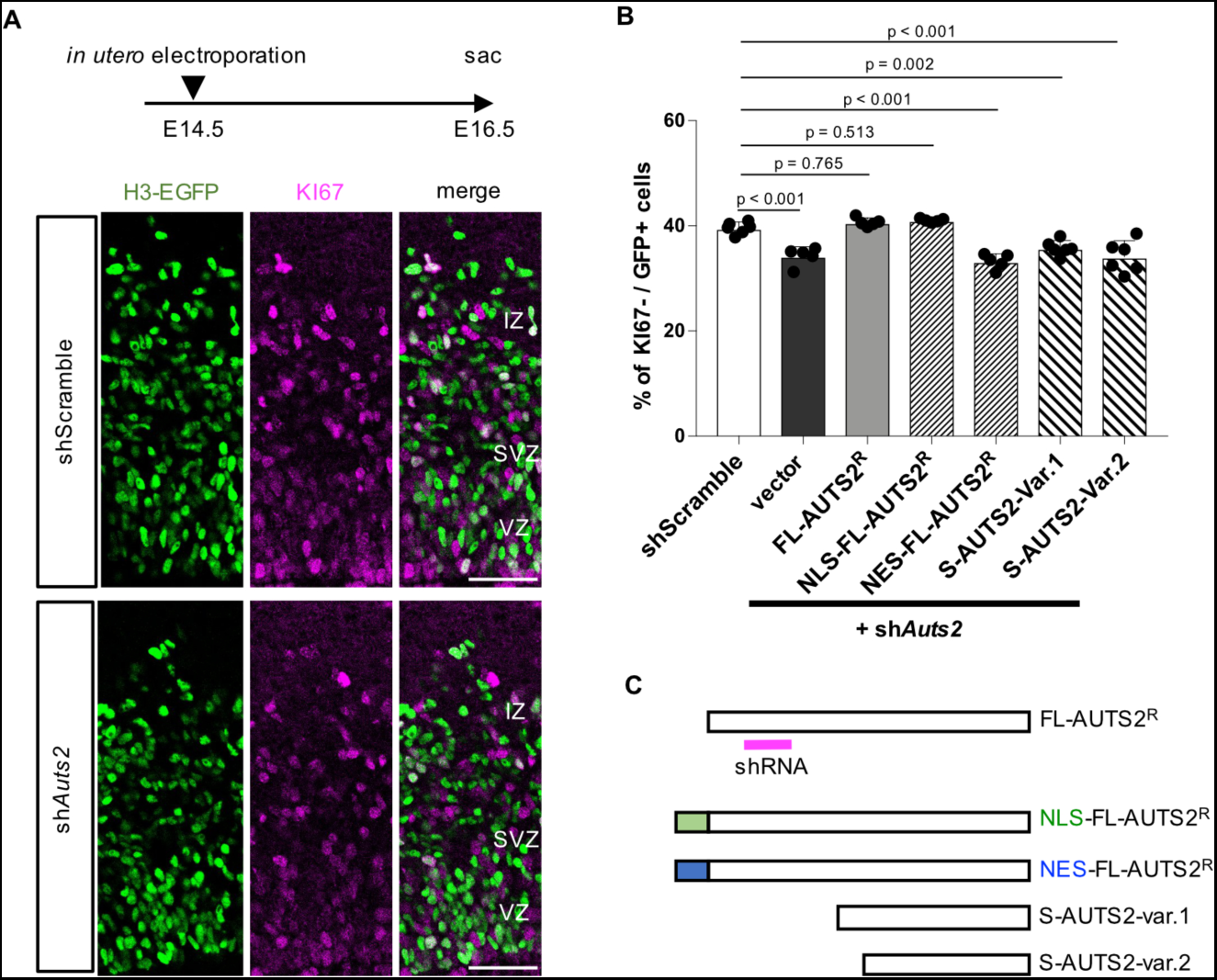
Nuclear AUTS2 is involved in neuronal production from neural progenitor cells. WT mouse embryos at E14.5 were co-electroporated with scrambled shRNA (shScramble) or *Auts2* shRNA (sh*Auts2*) together with empty or AUTS2 expression vectors indicated in (C) *in utero* and analyzed at E16.5. (A) Representative images of immunostaining for H3-EGFP (green) and KI67 (magenta) in the dorsolateral cerebral cortices of mice electroporated with shScramble and sh*Auts2* vectors. Scale bars, 50 µm. (B) Analyses of the ratio of KI67-negative cells to GFP^+^ cells. (C) Schematic diagrams of shRNA-resistant full-length-AUTS2 (FL-AUTS2^R^), NLS-FL-AUTS2^R^, NES-FL-AUTS2^R^, and C-terminal AUTS2 short variants (S-AUTS2-var.1 and var.2). The magenta bar indicates the shRNA binding site. NLS, nuclear localization signal; NES, nuclear export sequence. Data are presented as the mean ± SD (N = three mice, six sections); One-way ANOVA with Dunnett’s post-hoc test.

### FL-AUTS2 in nuclei is required for neuron production

Next, specific short hairpin RNA (shRNA) for *Auts2*^27^ or control scrambled shRNA vector was co-electroporated with a histone H3-fused green fluorescent protein (GFP) (H3-GFP) vector into the VZ of WT cortices at E14.5, followed by immunohistochemical analysis at E16.5. Under these experimental conditions in control, approximately 40% of the electroporated cells were KI67-negative cells, or putative postmitotic neurons produced from neural progenitors (Figures 3A and B). In contrast, introducing *Auts2* shRNA reduced this proportion, consistent with decreased neuron production due to prolonged cell cycle in *Auts2* KO IPCs. However, the reduced number of KI67-negative cells in *Auts2*-shRNA introduced cortex was fully rescued by the co-expression of shRNA-resistant full-length AUTS2 (FL-AUTS2^R^) (Figures 3 A and B). This rescuing effect was also observed for nuclear localization signal (NLS)-tagged FL-AUTS2^R^ (NLS-FL-AUTS2^R^)^10^ but not for the nuclear export sequence (NES)-tagged FL-AUTS2^R^ (NES-FL-AUTS2^R^)^27^ (Figures 3B and C). This suggests that nuclear-localizing AUTS2 is required for this developmental event. We also reported that AUTS2 short variants (var.1 and var.2) are expressed in the developing mouse cerebral cortex^27^. However, the short variants did not rescue the reduction in KI67-negative cells (Figures 3 B and C), indicating that these short variants were not involved in this event. These findings suggest that the nuclear localization of FL-AUTS2 is vital in neuronal production from progenitors.

### Loss of *Auts2* alters the expression of cell differentiation and cell proliferation-related genes

To isolate IPCs from the developing cerebral cortex by fluorescence-activated cell sorting (FACS), we newly generated a knock-in (KI) mouse line, in which a destabilized EGFP (d2EGFP) gene with a T2A self-cleaving peptide sequence was inserted into *Eomes* (coding TBR2 protein) genomic locus (Figures S4A–C). TBR2 and d2EGFP were bicistronically expressed in this mouse line in the IPCs. Indeed, we confirmed that the TBR2 and d2EGFP proteins were efficiently cleaved in cortical lysates from *Eomes^T2A-d2EGFP^* KI embryos at E14.5 (Figure S4D). It has been reported that *Eomes*-deficient homozygous mice showed embryonic lethality, reduced cortical size, and severe hypoplasia in olfactory bulbs^33^. In contrast, *Eomes^T2A-d2EGFP/T2A-d2EGFP^* homozygotes were born normally and exhibited a normal-sized cerebral cortex and olfactory bulb compared with WT mice (Figure S4E). We further observed that EGFP was exclusively expressed in TBR2-immunopositive cells in *the Eomes^T2A-d2EGFP/+^*cerebral cortex at E14.5 (Figure S4F), suggesting that IPCs were specifically labeled with EGFP without interfering with TBR2 protein expression and function in KI mice.

Using this mouse line, we tried to isolate IPCs and RGCs derived from the cerebral cortices of control (*Eomes^T2A-d2EGFP/+^; Auts2^flox/flox^*) or *Auts2* cKO homozygotes (*Eomes^T2A-d2EGFP/+^; Emx1^Cre/+^; Auts2^flox/flox^*) at E15.5, by FACS with EGFP-positivity (EGFP^+^ cells) and high expression of CD133 (PROM1, CD133^high^ cells), respectively (Figures 4A and B). We next performed RNA-sequencing (RNA-seq) for each isolated population. The expression of RGC marker genes, such as *Sox2*, *Nes*, *Pax6*, and *Hes1,* was enriched in CD133^high^ cells (Figure 4C), which were confirmed by the real-time quantitative PCR (RT-qPCR) analyses (Figure S4G). Immunocytochemistry revealed that most CD133^high^ cells expressed SOX2 but not TBR2 (Figure 4D). These observations indicated that most CD133^high^ cells were RGCs. In contrast, most EGFP^+^ cells were immunoreactive to TBR2 (Figure 4D). This suggests that cell sorting in *Eomes^T2A-d2EGFP/+^* KI mice was efficiently performed. In the RNA-seq and RT-qPCR analyses of EGFP^+^ cells, *Eomes* expression was concentrated, as expected (Figures 4C and S4G). In the EGFP^+^ cell population, expression of immature neuronal markers, such as *NeuroD1* and *Tubb3*, was detected compared to CD133^high^ cells (Figures 4C and S4G). This may reflect the small population (15%) of postmitotic cells in TBR2^+^ cells (Figure S3B), consistent with reports that TBR2^+^ cells contain a small proportion of postmitotic immature neurons^31,34^. These findings suggest that the EGFP^+^ cell population mainly includes IPCs. *Auts2* expression was more enriched in EGFP^+^ cells than in CD133^high^ cells (Figure 4C), suggesting that *Auts2* expression was higher in IPCs than in RGCs.

**Figure 4:**
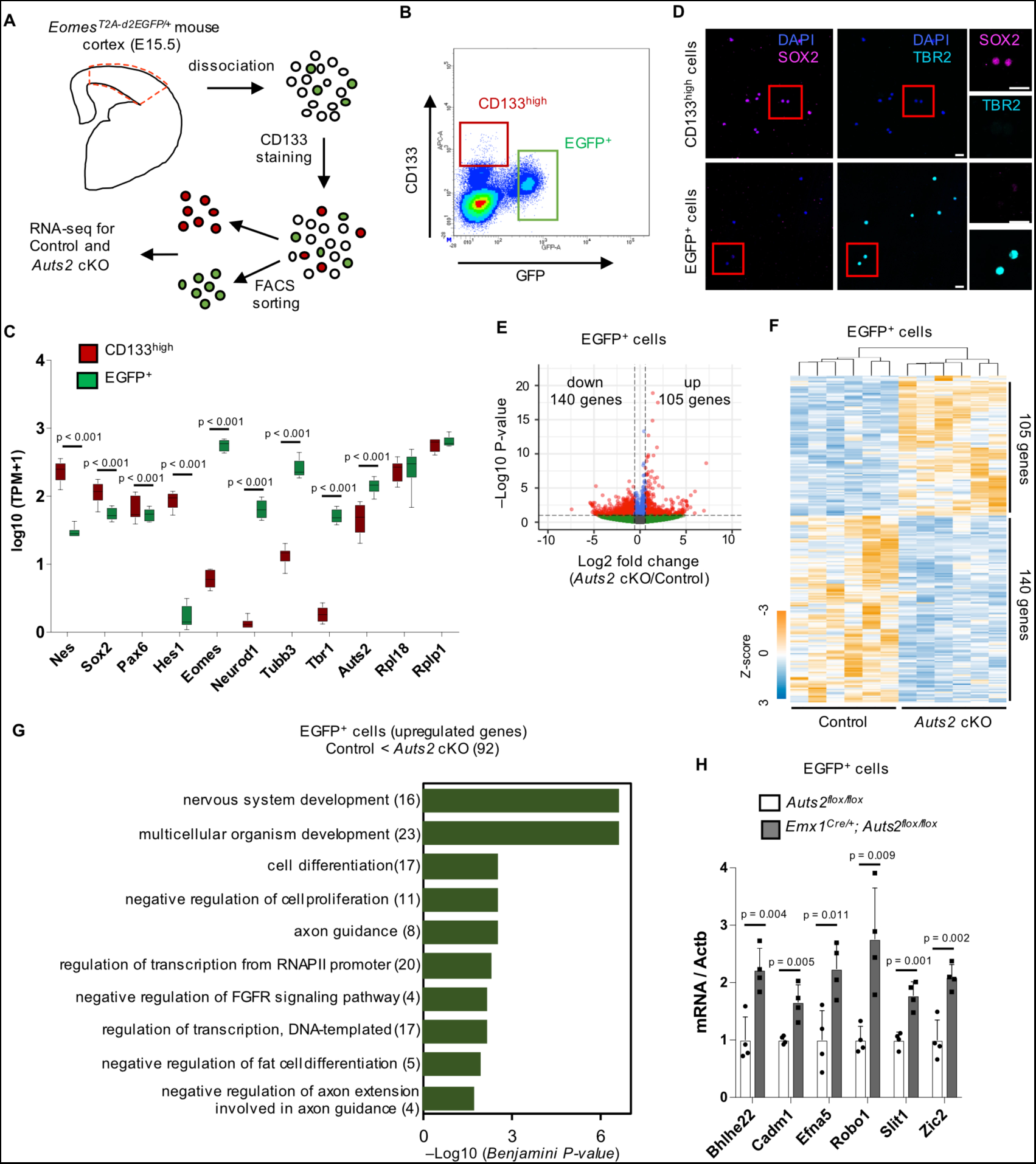
Transcriptional profiling of IPCs in control and *Auts2* cKO mice at E15.5. (A) Experimental design for transcriptional analysis. CD133^high^ and EGFP^+^ cells were sorted from *Eomes^T2A-d2EGFP/+^; Auts2^flox/flox^*(Control) or *Eomes^T2A-d2EGFP/+^; Emx1^Cre/+^; Auts2^flox/flox^*(*Auts2* cKO) cortices at E15.5. (B) Representative plot showing sorting gates for CD133^high^ and EGFP^+^ cells. (C) Expression levels of *Nes, Sox2, Pax6, Hes1, Eomes, Neurod1, Tubb3, Tbr1, Fezf2, Auts2, Rpl18,* and *Rplp1* transcripts in sorted each cell type from E15.5 control mouse cortex. *Rpl18* and *Rplp1* are presented as housekeeping genes. TPM, Transcripts per million. *Adjusted* P-value calculated by DESeq2 was shown. (D) Immunostaining of CD133^high^ and EGFP^+^ cells sorted by FACS from E15.5 *Eomes^T2A-d2EGFP/+^* cortex with DAPI (blue), anti-SOX2 (magenta) and anti-TBR2 (cyan) antibodies. Scale bars, 20 µm. (E) Volcano plot showing differences in gene expression in EGFP^+^ cells between control and *Auts2* cKO mice. Red dots indicate upregulated and downregulated genes (P-value < 0.01 and |Log_2_ fold-change| > 0.58). (F) Heatmap for the differential expressed genes (DEG) RNA levels in EGFP^+^ cells from control and *Auts2* cKO mice. The color scale is shown on the left bottom. (G) DAVID Gene Ontology biological process analysis of upregulated genes in EGFP^+^ cells. The top 10 GO terms with Benjamini-Hochberg adjusted P-value are shown. Numbers in parentheses indicate the gene count. (H) RT-qPCR analysis for *Bhlhe22, Cadm1, Efna5, Robo1, Slit1, and Zic2* in sorted EGFP^+^ cells from control and *Auts2* cKO mice at E15.5. Data are presented as the mean ± SD (N=four mice); unpaired Student’s t-test. All transcriptome analyses used six biological replicates per genotype.

In RNA-seq analysis, we identified 8619 differentially expressed genes (DEGs) between CD133^high^ cells (mostly RGCs) and EGFP^+^ cells (mainly IPCs) (Figure S5A), which may reflect the different cell characters. In EGFP^+^ cells, we found that the expression of 105 genes was significantly upregulated, and that of 140 genes was downregulated in cKO homozygotes compared with the control (Figures 4E and F). We named these genes “IPC-upregulated-genes” and “IPC-downregulated-genes,” respectively.

Gene Ontology (GO) term analysis demonstrated that IPC-upregulated-genes were associated with biological processes, including “nervous system development,” “cell differentiation,” and “negative regulation of cell proliferation” (Figure 4G), implying that AUTS2 functions in IPCs to regulate proliferation and/or differentiation. We performed RT-qPCR using EGFP^+^ cells for some of the IPC-upregulated-genes, confirming that their expression was upregulated in the cKO compared with that in the control (Figure 4H). There was no significant GO term for the IPC-downregulated-genes. We also identified DEGs in sorted CD133^high^ cells between control and cKO mice. The expression of 192 genes was significantly upregulated, whereas that of 190 genes was downregulated (Figure S5B), which gave one GO term (Figure S5C). Notably, the DEGs for CD133^high^ and EGFP^+^ cells did not overlap very much (Figure S5D), suggesting that AUTS2 might control the expression of different genes in RGCs and IPCs.

### AUTS2 is involved in IPC proliferation through suppressing *Robo1* expression

As we observed a significant phenotype in IPCs but not in RGCs, we decided to focus on IPCs. Among IPC-upregulated- and IPC-downregulated-genes, we investigated the *Robo1* gene, which was also associated with the GO terms “nervous system development” and “negative regulation of cell proliferation” (Figure 4G). Expression of this gene increased approximately 2.5-fold in *Auts2* cKO EGFP^+^ cells compared with that in the control (Figure 4H). We electroporated the overexpression vector (OE) for *Robo1* into the VZ of WT cortices at E13.5 and analyzed them at E15.5 when the reduction in IPC division was observed in the *Auts2* mutant (Figure 2B). We found that *Robo1* OE decreased the ratio of mitotic cells (PH3^+^) among electroporated TBR2^+^ cells compared with the control (Figure 5A). In addition, *Robo1* OE at E14.5 resulted in reduced postmitotic (KI67-negative and GFP^+^) cells at E16.5 (Figure 5B), consistent with our observation that electroporation of the *Auts2* KD vector decreased the proportion of postmitotic cells at 2 days after electroporation (Figures 3A and B). These findings suggest that *Robo1* can suppress the proliferation of IPCs and the production of an appropriate number of neurons.

**Figure 5:**
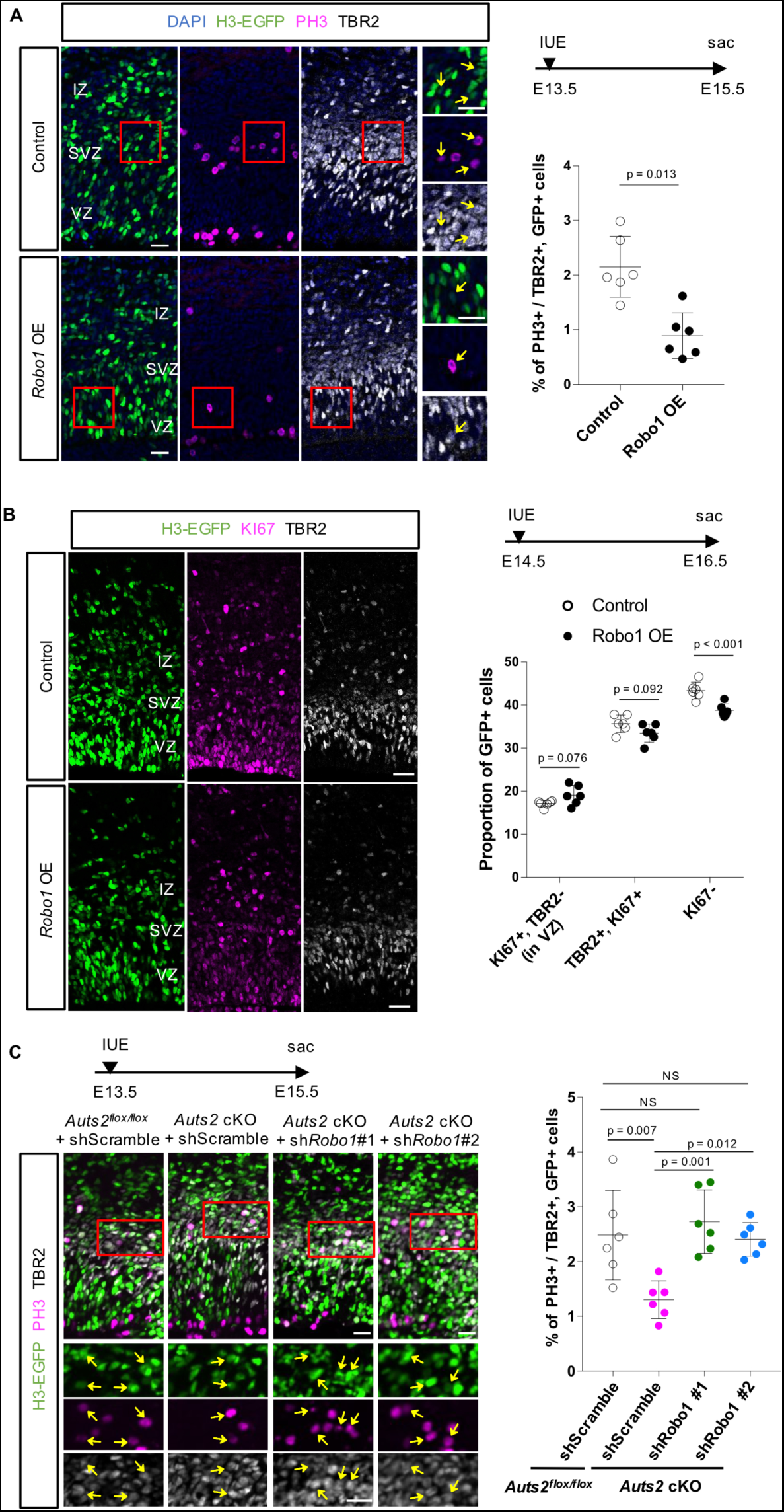
Elevated expression of *Robo1* leads to a decrease in IPC proliferation. (A) *In utero* electroporation (IUE) of either an empty vector (control) or a *Robo1* expression vector (*Robo1* OE) together with a H3-GFP expression vector was performed into WT cortices at E13.5. The cortical sections are stained with DAPI (blue), anti-GFP (green), anti-PH3 (magenta), and anti-TBR2 (white) antibodies at E15.5. The graph shows the ratio of PH3^+^ cells in TBR2^+^ and GFP^+^ cells. Arrows indicate GFP^+^, PH3^+^, and TBR2^+^ cells. (B) IUE of an empty (control) or a *Robo1* expression vector (*Robo1* OE) and a H3-GFP expression vector into WT cortices at E14.5. The cortical sections are immunostained with anti-GFP (green), anti-KI67 (magenta), and anti-TBR2 (white) antibodies at E16.5. The graph shows the proportion of KI67^+^ and TBR2-negative cells located in the VZ (RGCs), TBR2^+^ and KI67^+^ cells (IPCs), and KI67-negative differentiated cells (postmitotic neurons) among total GFP^+^ cells. The percentage of RGCs and IPCs in electroporated cells was not different between *Robo1* OE and control. (C) *Auts2^flox/flox^* or *Emx1^Cre/+^; Auts2^flox/flox^*(*Auts2* cKO) cortices were electroporated with scrambled or *Robo1* shRNA vector *in utero* at E13.5 and immunostained with anti-GFP (green), anti-PH3 (magenta) and anti-TBR2 (white) antibodies at E15.5. The graph shows the percentage of PH3^+^ cells among TBR2^+^ and GFP^+^ cells. Arrows indicate GFP^+^, PH3^+^, and TBR2^+^ cells. Data are presented as the mean ± SD (N = three mice, six sections). NS, not significant, unpaired Student’s t-test (A, B) and One-way ANOVA with Turkey’s post-hoc test (C). Scale bars, 20 µm (A, C) and 50 µm (B).

We performed a rescue experiment by *in utero* electroporation of *Robo1* KD vectors into *Auts2* cKO cortices to test whether the reduced IPC proliferation and neuron production in *Auts2* KO mice was caused by increased expression of *Robo1*. We generated two *Robo1*-specific shRNAs that effectively depleted ROBO1 protein (Figure S6A). Each *Robo1* shRNA or scrambled shRNA was electroporated into the VZ of *Auts2* cKO homozygous cortices at E13.5, and immunostaining for TBR2 and PH3 was performed 2 days after electroporation (E15.5). We found that the ratio of mitotic cells in electroporated TBR2^+^ cells in *Auts2* cKO cortices was efficiently recovered by electroporation with either *Robo1* shRNA to the same degree as in the scrambled shRNA-transfected control (*Auts2^flox/flox^*) cortices (Figure 5C). Next, we performed another rescue experiment in which each *Robo1* shRNA or scrambled shRNA was electroporated into the VZ of *Auts2* cKO homozygous cortices at E14.5, and immunostaining for KI67 and TBR2 was performed at E16.5. Among the electroporated cells, the reduction in KI67-negative differentiated cells in *Auts2* cKO cortices was efficiently rescued by introducing either *Robo1* shRNA (Figure S6B). These findings suggest that AUTS2 promotes the proliferation of IPCs and the production of an appropriate number of neurons by repressing *Robo1* expression.

### AUTS2 represses the expression of genes related to neuronal differentiation in IPCs by maintaining repressive chromatin status

AUTS2 has been reported to regulate gene expression in the cell nuclei^9,12,13,28,35^; therefore, we next tried to identify AUTS2-interacting genomic regions in IPCs by conducting a cleavage under targets and tagmentation (CUT&Tag) experiment^36^ to the sorted EGFP^+^ cells from control (*Eomes^T2A-d2EGFP/+^; Auts2^flox/flox^)* and cKO (*Eomes^T2A-d2EGFP/+^; Emx1^Cre/+^; Auts2^flox/flox^)* cortices at E15.5 using the anti-AUTS2 antibody, followed by deep-sequencing. In the control cells, we identified multiple genomic loci to which AUTS2 bound significantly and strongly (Figure 6A). In contrast, global levels of AUTS2 binding were reduced in *Auts2* cKO homozygotes (Figures 6A and S7A), suggesting that the CUT&Tag experiment worked efficiently. However, the signals in the cKO cells were not completely lost. A truncated isoform (S-AUTS2-var.2) is expressed even in *Auts2* cKO cells^27^, and the AUTS2 antibody can recognize that isoform; therefore, the signals for cKO might correspond to loci bound to S-AUTS2-var.2. In this experiment, we successfully identified 5464 significant AUTS2-binding loci in control cells.

**Figure 6:**
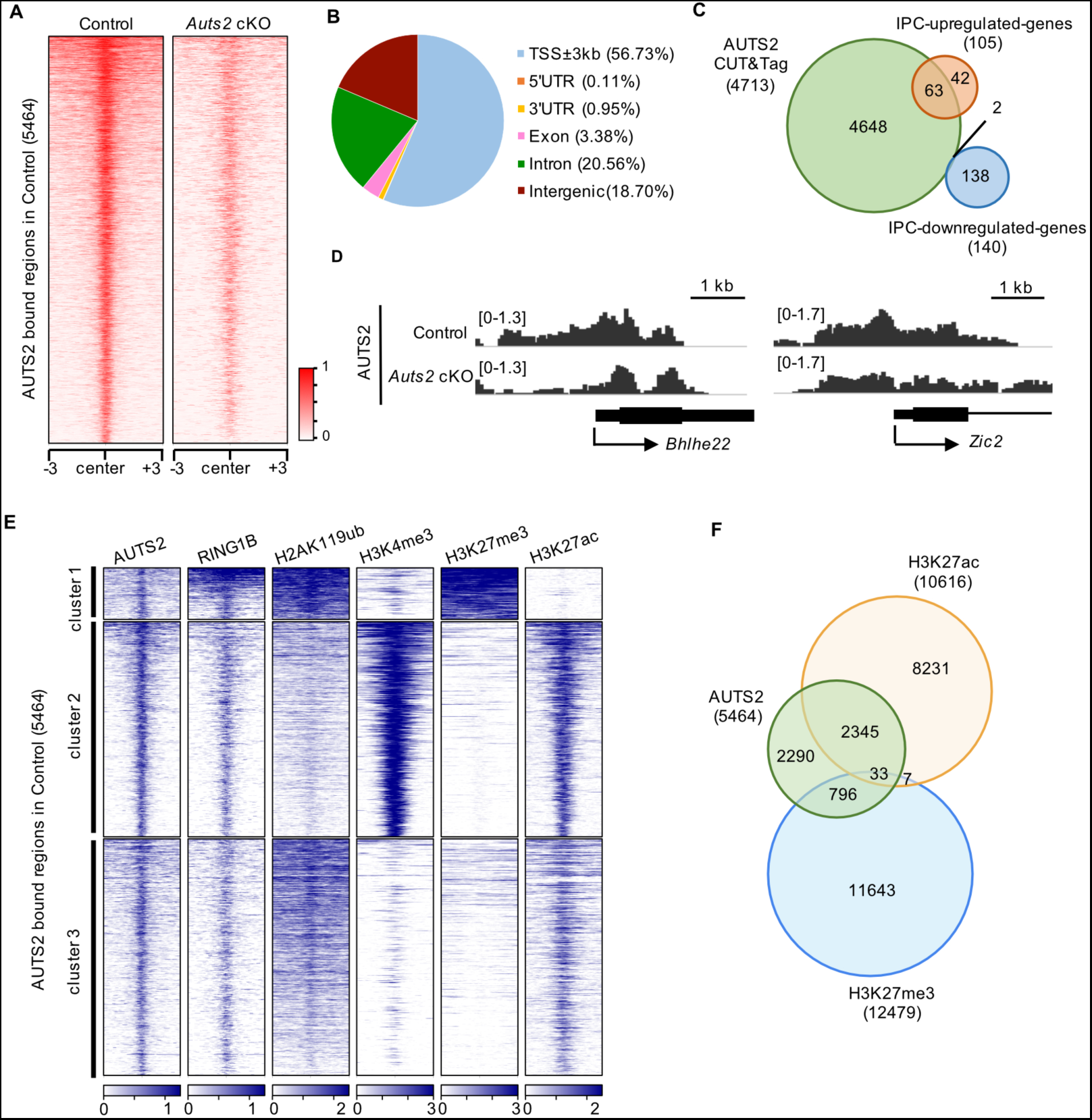
AUTS2 binds to the active chromatin and the repressive chromatin. (A) Heatmap showing the CUT&Tag signals of AUTS2 in control (left) and *Auts2* cKO homozygous (right) cells centered on AUTS2-binding loci (±3 kb) identified in control at E15.5. (B) Pie chart showing the percentage of AUTS2-binding loci on each genomic region. (C) Genes near the AUTS2-binding loci were identified using the Genomic Regions Enrichment of Annotations Tool (GREAT). Venn diagrams show the overlap between those genes and IPC-upregulated or IPC-downregulated genes identified by RNA-seq. (D) Interactive Genomics Viewer (IGV) browser views showing the CUT&Tag signal for AUTS2 in control and *Auts2* cKO cells at the indicated loci. (E) K-means clustering of AUTS2, RING1B, H2AK119ub, H3K4me3, H3K27me3, and H3K27ac CUT&Tag signals in control cells centered on AUTS2 binding peaks (±3 kb) identified in control. (F) Venn diagrams showing the extent of overlap for AUTS2-binding loci, H3K27ac-and H3K27me3-modified loci.

AUTS2-binding loci were predominantly detected near or within genes (81.30%), especially the ± 3 kb region around transcriptional start sites (TSS) (56.73%) and occasionally in introns, exons and untranslated regions (UTRs). However, only 18.70% was found in intergenic regions (Figure 6B). We identified 4713 genes in the vicinity of AUTS2 binding loci, which included 63 of 105 IPC-upregulated-genes (60%) but only 2 of 140 IPC-downregulated-genes (1.4%) (Figure 6C). Previous studies have shown that AUTS2 is involved in transcriptional activation and repression in a cell type-dependent manner^28^. These findings suggest that AUTS2 might be involved in suppressing gene transcription in IPCs and that the 63 genes, which included *Robo1*, might be direct target candidates of AUTS2. Among them, *Bhlhe22* and *Zic2* were representative genes in which AUTS2 binding signals were abundant around the TSS in the control but were reduced in *Auts2* cKO cells (Figure 6D). A similar tendency was also observed for the binding of AUTS2 to the region around the TSS of the *Robo1* gene (Figure 7D).

**Figure 7:**
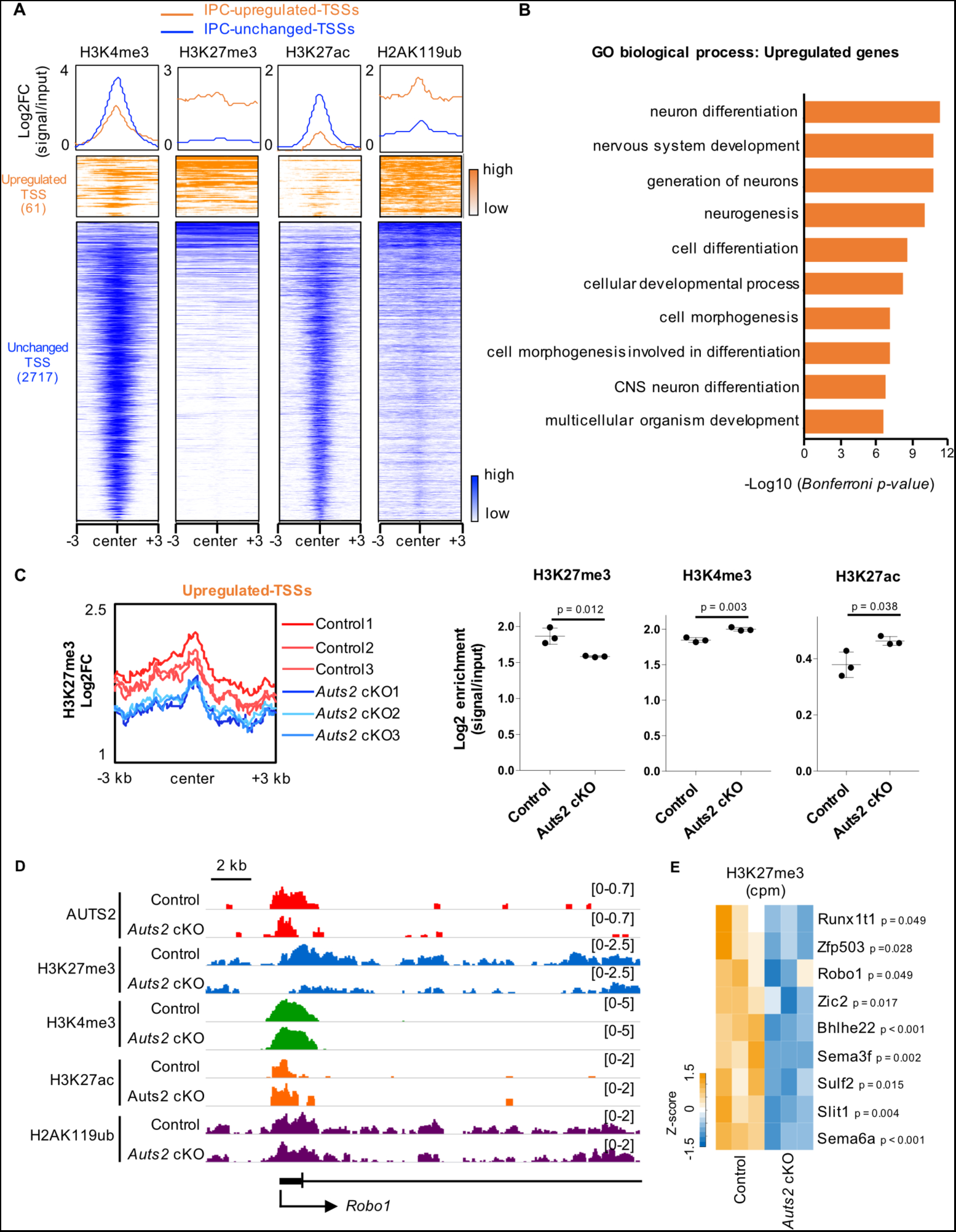
AUTS2 maintains the repressive chromatin status of genes related to neuronal differentiation. (A) Density plots and heatmap of H3K4me3, H3K27me3, H3K27ac, and H2AK119ub CUT&Tag signals centered on AUTS2-binding loci (±3 kb) localized near IPC-upregulated-TSSs (orange) and IPC-unchanged-TSSs (blue). FC: fold change. (B) Graph showing the top 10 GO-term biological processes for genes with IPC-upregulated-TSSs on GREAT. (C) Density profile shows H3K27me3 occupancy in control and *Auts2* cKO cells centered on IPC-upregulated-TSSs (±3 kb). Graphs showing the enrichment of H3K27me3, H3K4me3, and H3K27ac on IPC-upregulated-TSSs. FC, fold change. N = three biological replicates. (D) IGV browser views showing the CUT&Tag signals for AUTS2, H3K27me3, H3K4me3, H3K27ac and H2AK119ub in control and *Auts2* cKO cells at *Robo1* locus. H3K4me3 and H3K27ac levels were significantly increased (H3K4me3, p = 0.003; H3K27ac, p = 0.039). H2AK119ub level was not significant (p = 0.182). N = three biological replicates, unpaired Student’s t-test. (E) Heatmap of count per million (cpm) for H3K27me3 on AUTS2-binding loci located near the representative IPC-upregulated-genes related to “neuron differentiation (GO:0030182)” and/or “cell differentiation (GO:0030154).” Data are presented as the mean ± SD; unpaired Student’s t-test.

As previous reports have shown that AUTS2 interacts with PRC1.3/5 complexes^9,13,28^, we performed CUT&Tag experiments on RING1B, a core subunit of PRC1, in sorted EGFP^+^ cells from control mice (*Eomes^T2A-d2EGFP/+^; Auts2^flox/flox^*) at E15.5. Approximately 60% of AUTS2-binding loci were also RING1B-binding loci (Figure S7B). We conducted CUT&Tag experiments for H2AK119ub, H3K4me3, H3K27me3, and H3K27ac in the same cells to investigate the histone modifications around the AUTS2-binding loci in IPCs. K-means clustering of the data for AUTS2, RING1B, H2AK119ub, H3K4me3, H3K27me3, and H3K27ac showed that AUTS2-binding loci could be classified into three clusters (Figure 6E). Cluster 1 loci were suggested to correspond to repressive chromatin because they were enriched with H3K27me3 and H2AK119ub but not with H3K4me3 or H3K27ac. In contrast, cluster 2 loci may correspond to active chromatin because they are enriched with H3K4me3 and H3K27ac but not with H3K27me3 or H2AK119ub. Previously, it was shown that AUTS2-binding loci were predominantly localized in active chromatin in experiments using P1 whole-brain and mouse embryonic stem cell (mESC)-derived motor neurons^9,13^. Notably, our experiments showed that AUTS2-binding loci were present on active chromatin (cluster 2) and repressive chromatin (cluster 1), at least in IPCs (Figure 6E). In this experiment using control IPCs, we identified 12479 and 10616 significant loci that are histone modified in H3K27me3 and H3K27ac, respectively. There was significant overlap between both types of loci and the AUTS2-binding loci, suggesting that AUTS2 binds to both active and repressive chromatin (Figure 6F).

Next, we focused on the AUTS2-binding loci within 3 kb of the TSS of IPC-upregulated-genes (61 loci) because more than half of the AUTS2-binding loci were localized within 3 kb of TSSs (Figure 6B). As a control, we also identified 2717 AUTS2-binding loci within 3 kb of the TSS of genes whose expression did not change in the cKO IPCs (P-value > 0.1). We named these loci as “IPC-upregulated-TSSs” and “IPC-unchanged-TSSs” (Figure 7A). We compared histone modifications between IPC-upregulated-TSSs and IPC-unchanged-TSSs in the control genetic background (*Eomes^T2A-d2EGFP/+^; Auts2^flox/flox^*). IPC-upregulated-TSSs were enriched with repressive histone marks (H3K27me3 and H2AK119ub), whereas they harbored poorly active histone marks (H3K4me3 and H3K27ac) compared with IPC-unchanged-TSSs (Figures 7A and S8A). This suggests that the chromatin of IPC-upregulated-TSSs might be relatively in the repressive condition. In addition, GO analysis in biological processes showed that the genes with IPC-upregulated-TSSs were associated with the GO terms, including “neuron differentiation,” “generation of neurons,” “neurogenesis,” and “central nervous system (CNS) neuron differentiation” in contrast to GO terms for the genes with IPC-unchanged-TSSs (Figures 7B and S8B). These findings suggest that AUTS2 suppresses the expression of genes associated with neuron production in IPCs.

We further conducted the CUT&Tag experiments on sorted EGFP^+^ cells from *Auts2* cKO mice at E15.5, using antibodies for histone modifications and RING1B to test whether histone modifications and RING1B-binding were affected without the AUTS2 protein. The levels of RING1B binding to IPC-upregulated-TSSs and IPC-unchanged-TSSs did not significantly change in cKO cells compared with control cells (Figure S8C), suggesting that AUTS2 is not involved in the localization of the PRC1 complex on the chromatin in IPCs. H2AK119ub modification was not affected in either IPC-upregulated- or unchanged-TSSs (Figure S8D), indicating that AUTS2 may not play a major role in chromatin ubiquitination, at least in IPCs. Regarding IPC-upregulated-TSSs, the levels of the repressive modification (H3K27me3) were reduced in cKO cells, whereas those of the active modifications (H3K4me3 and H3K27ac) were elevated (Figures 7C and S8E). However, these tendencies were not observed for IPC-unchanged-TSSs (Figure S8F). At the TSS of the *Robo1* gene, a representative IPC-upregulated-gene, the repressive mark (H3K27me3) was reduced, and active marks were slightly but significantly increased in the mutant cells compared with the control (Figure 7D). Similarly, a significant reduction of the H3K27me3 modification was observed in other representative genes with IPC-upregulated-TSSs, associated with the GO terms “neuron differentiation,” “generation of neuron,” and/or “cell differentiation” (Figure 7E). These findings suggest that AUTS2 maintains the repressive states of genes associated with neuron production in IPCs.

### AUTS2 interacts with PRC2 proteins in IPCs

H3K27me3 modification is catalyzed by PRC2^37–39^. We performed co-immunoprecipitation experiments to investigate whether AUTS2 interacts with PRC2. First, to efficiently immunoprecipitate sufficient amounts of exogenous AUTS2 protein, 3xFlag-tagged FL-AUTS2 (FLAG-AUTS2) was transfected into Neuro2a cells and immunoprecipitated using an anti-FLAG antibody 2 days after transfection. Endogenous EZH2 and SUZ12, the core proteins of PRC2, were found to be co-immunoprecipitated only in FLAG-AUTS2 expressing cells (Fiure 8A). Next, we immunoprecipitated endogenous AUTS2 proteins with an anti-AUTS2 antibody from E14.5 cerebral cortices. Western blotting showed that EZH2 and SUZ12 were co-immunoprecipitated with AUTS2 (Figure 8B), suggesting that AUTS2 may interact with PRC2 in the developing cerebral cortex. Furthermore, we performed a proximity ligation assay (PLA) using anti-AUTS2 and anti-EZH2 antibodies in the sorted EGFP^+^ cells from the *Eomes^T2A-d2EGFP/+^* cerebral cortices at E15.5. Notably, many puncta were observed in the cell nuclei with both antibodies compared with those with a single antibody (Figure 8C), which also suggested that AUTS2 interacts with PRC2 in the IPC nuclei.

**Figure 8:**
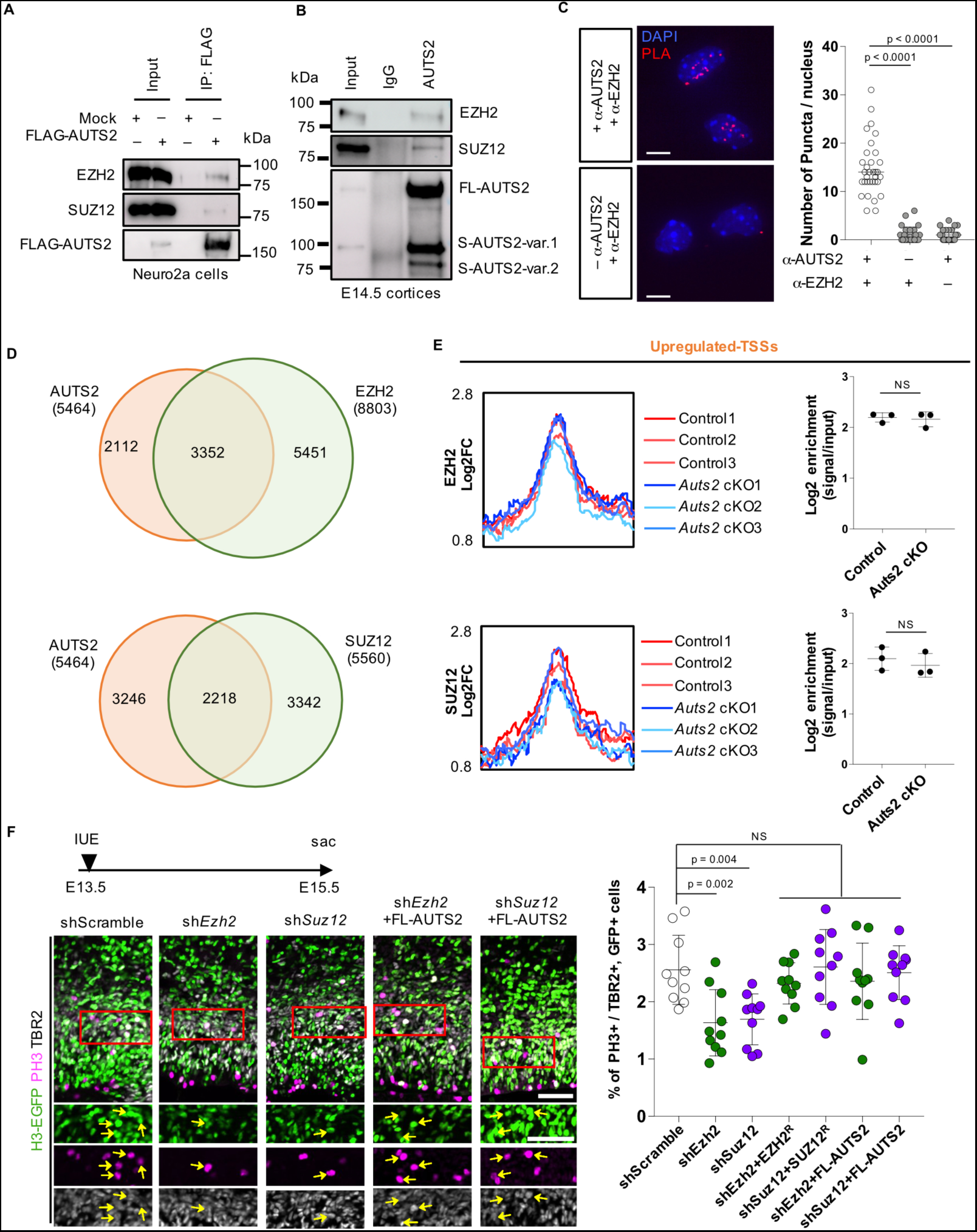
AUTS2 interacts with PRC2 in IPCs. (A) Co-immunoprecipitation assay using Neuro2a cells. 3xFLAG-tagged FL-AUTS2 expression vector or 3xFlag-tagged empty vector was transfected into Neuro2a cells. The nuclear lysates were incubated with an anti-FLAG antibody. Western blotting was performed with anti-EZH, anti-SUZ12, and anti-FLAG antibodies. (B) Co-immunoprecipitation assay using E14.5 cerebral cortices. Rabbit anti-AUTS2 and rabbit control IgG antibodies were used for immunoprecipitation. The precipitates were immunoblotted with anti-EZH2, anti-SUZ12, and anti-AUTS2 antibodies. (C) Proximity ligation assay (PLA) with rabbit anti-AUTS2 and mouse anti-EZH2 antibodies in sorted EGFP^+^ cells from E15.5 WT cortices. Representative images show the PLA signals (red) and DAPI (blue) in cells incubated with both antibodies (top) or a single antibody (only anti-EZH2, bottom). The graph shows the number of puncta in the nucleus of each sample. N = three biological replicates, 28–35 cells. (D) Venn diagrams showing the extent of overlap between AUTS2 peaks and EZH2 (top) or SUZ12 (bottom) peaks. (E) Density profiles show the occupancy of EZH2 (top) and SUZ12 (bottom) in control and *Auts2* cKO cells centered on IPC-upregulated-TSSs (±3 kb). Graphs show the enrichment of EZH2 and SUZ12 on IPC-upregulated-TSSs. N = three biological replicates. (F) IUE of indicated vectors and an H3-GFP expression vector were performed into WT cortices at E13.5. The cortical sections are stained for GFP (green), PH3 (magenta), and anti-TBR2 (white) antibodies at E15.5. The graph shows the ratio of PH3^+^ cells in TBR2^+^ and GFP^+^ cells. Red rectangles indicate the location of the images below. Arrowheads indicate GFP^+^, PH3^+^, and TBR2^+^ cells. N = five mice, 10 sections. Data are presented as the median (C) and the mean ± SD (E, F). NS, not significant, Kruskal–Wallis test (C), unpaired Student’s t-test (E), and One-way ANOVA with Dunnett’s post-hoc test (F). Scale bars, 5 µm (C), 50 µm (F).

We performed CUT&Tag experiments for EZH2 and SUZ12 on sorted EGFP^+^ cells from control cortices at E15.5 to examine the correlation of PRC2-binding loci with AUTS2 ones. Global distributions of EZH2 and SUZ12 were observed at AUTS2-binding loci (Figure S9A), and 3352 and 2218 AUTS2 peaks overlapped with EZH2 and SUZ12 peaks, respectively (Figure 8D). These findings suggest that AUTS2 co-localizes with PRC2 in the genome of IPCs. We further conducted CUT&Tag experiments in the sorted EGFP^+^ cells from *Auts2* cKO cortices. Then, we compared the levels of EZH2- and SUZ12-binding to IPC-upregulated-TSSs between control and *Auts2* cKO cells to investigate whether PRC2 localization was dependent on AUTS2. We did not detect differences in the binding levels to IPC-upregulated-TSSs or IPC-unchanged-TSSs (Figures 8E and S9B). Consistently, EZH2- and SUZ12-binding levels were not affected around the TSS of *Robo1* (Figure S9C). These findings suggest that AUTS2 is not involved in the localization of PRC2 on chromatins.

We performed KD experiments using shRNAs for *Ezh2* and *Suz12* to test the involvement of PRC2 in IPC division *in vivo*. First, we confirmed that the shRNA vectors sufficiently reduced the exogenous expression of *Ezh2* and *Suz12* in HEK293T cells (Figure S9D). These shRNA vectors plus the H3-GFP vector were co-electroporated into the VZ of the WT cortex at E13.5, and the sections were immunostained with PH3, TBR2, and GFP at E15.5. The ratio of PH3^+^ cells in TBR2^+^ and GFP^+^ cells was significantly reduced by sh*Ezh2* or sh*Suz12*, which was rescued by the co-introduction of shRNA-resistant EZH2 or SUZ12 (EZH2^R^ and SUZ12^R^) (Figure 8F). This suggests that PRC2 is required for the division of IPCs in the embryonic cortices. Interestingly, the co-introduction of FL-AUTS2 with *Ezh2* or *Suz12* shRNA vector restored the ratio of PH3^+^ cells (Figure 8F), implying that AUTS2 and PRC2 may work together in the same pathway for IPC division.

## Discussion

Previously, AUTS2 syndrome has been reported to present with a high frequency of microcephaly in addition to psychiatric disorders such as ID and ASD^4,6,7^. A previous study demonstrated that a patient with the *AUTS2^T534P^* missense mutation displayed reduced cortical area and ventriculomegaly on MRI and that a human cerebral organoid study indicated that the *AUTS2^T534P^* missense mutation impairs organoid growth^8^. In its mutant organoids, the division of neural progenitor cells, presumably apical RGCs, is reduced compared to controls during the early neurogenic phase^8^. These suggest that cerebral organoids with *AUTS2* missense mutation can reproduce the microcephaly of the AUTS2 syndrome and that AUTS2 may have some roles in neurogenesis. However, the mechanism by which microcephaly is caused by loss of *AUTS2* remains unclear, and microcephaly-like phenotypes have not been reproduced in *in vivo* models of animals with neocortex. In this study, we demonstrated that *Auts2* cKO mice exhibit a reduced number of cortical upper-layer neurons and reduced cortical thickness (Figures 1C–E). Notably, this phenotype was observed in *Auts2* heterozygous cKO mice, which may mimic the pathology of human AUTS2 syndrome patients. In *AUTS2* syndrome, different genomic mutations lead to a variety of clinical features and symptoms, but not all symptoms are present in individual patients ^2,4^. Approximately 65% of patients with *AUTS2* syndrome exhibit microcephaly^4^. Symptomatic diversity is believed to be caused by different mutations at various sites in the very long *AUTS2* gene. Correspondingly, a recent study reported that *Auts2* KO mice with an allele that differs from *Auts2^del8^* did not show any change in the number of neurons in the cortex^12^. So far, several mouse lines with different mutations in the *AUTS2* gene have been reported^10,12,13,27,40^. These mouse models may serve as models for *AUTS2* syndrome with various genomic variants.

Microcephaly is often caused by a reduction in the number of cortical neurons. During evolution, mammals are thought to have acquired IPCs in addition to RGCs, thereby leading to an increase in the number of neurons and overall size of the cerebral cortex ^24^. Moreover, during the evolution to primates, including humans, the acquisition of outer radial glial cells (oRGCs) has led to a further increase in the number of neurons and expansion of the cerebral cortex^41,42^. Unfortunately, because oRGCs are present in very limited numbers in mice^26^, analyses using mouse models allow us to study the effects of the *Auts2* mutations on RGCs and IPs, but not to observe the effects on oRGCs. Nevertheless, our *Auts2* cKO mice (*Auts2^del8^/Emx1^Cre^*) may recapitulate at least some of the pathophysiology of microcephaly in human AUTS2 syndrome, since we observed a decrease in dividing IPCs and a decrease in the number of upper layer neurons and thickness of the cerebral cortex.

Previous clonal analyses have shown that mouse IPCs divide once to multiple times, generating approximately 2∼12 neurons^19,21^. However, the regulatory mechanisms of their proliferation and differentiation are still largely unexplored compared to RGCs. We found that the protein ROBO1 suppresses IPC division (Figure 5A). AUTS2, in turn, suppresses *Robo1* expression in IPCs, leading to an increase in IPC division and the production of upper-layer neurons. Notably, the division of TBR2^+^ cells is absent or rare in reptiles or birds but substantially present in mammals, with increased IPC division in primates^43–47^. Therefore, this study provides insights not only into the regulatory mechanisms of IPC division and neuronal differentiation but also into the expansion of the mammalian brain during evolution. From the pathological point of view, we show that loss of *Auts2* expression increases *Robo1* expression in IPCs, elongates the cell cycle of IPCs, suppresses IPC divisions, and decreases the number of upper-layer neurons. This may explain, at least in part, the pathomechanisms of microcephaly of AUTS2 syndrome. The contribution of oRGCs to microcephaly in AUTS2 syndrome may have to wait for future studies, which may include long-term organoid culturing using primate cells. Through RNA-seq analysis in *Auts2* cKO mice, we were able to identify hundreds of DEGs in RGCs (Figure S5B). Although our histological analysis did not reveal any differences in RGC proliferation (Figure 2A), AUTS2 may still have a role to play in RGCs.

Previous studies have revealed that nuclear AUTS2 can function as a transcriptional activator or repressor, depending on the type of cell^28^. Using a luciferase reporter with UAS sequence, Monderer-Rothkoff et al showed that GAL4-AUTS2 expression increased luciferase activity in HEK293 cells but decreased in Neuro2a cells^28^. This indicates that AUTS2 might have a role in modifying histones to either activate or deactivate transcription, depending on the specific cell type. Gao et al. showed that, in HEK293 cells, AUTS2 interacts with non-canonical PRC1 and contributes to transcriptional activation by recruiting p300 and engaging in acetylation of H3K27^13^. In this study, we show that the AUTS2 protein is primarily involved in transcriptional repression in IPCs. In the *Auts2* cKO IPCs, we found many upregulated and downregulated genes, but among them, the AUTS2 protein binds mostly to the genomic regions around the TSSs of the upregulated genes (Figures 6B and C). Around the TSSs of those genes, histone modification of H3K27me3 was significantly reduced in the *Auts2* cKO IPCs (Figure 7C). We further found that AUTS2 interacts with EZH2 and SUZ12, the core components of PRC2 that catalyze the trimethylation of H3K27 (Figures 8A–C)^37–39^. In both CUT&Tag experiment and PLA, we showed that AUTS2 and PRC2 components (EZH2 and SUZ12) bind to common genomic regions in IPCs (Figures 8C, D and S9A). Electroporation of a shRNA vector for *Ezh2* or *Suz12* into the embryonic VZ decreased dividing IPCs, which was rescued by overexpression of AUTS2 (Figure 8F). These findings suggest that PRC2 and AUTS2 are cooperatively involved in the regulation of IPC divisions by trimethylating H3K27 and repressing gene expression. Regarding the *Robo1* gene, we found that PRC2 binds the region around the TSS in the control IPCs (Figure S9C). However, that region showed reduced levels of H3K27me3 in the cKO condition (Figure 7D). Consistently, previous studies have reported that *Robo1* expression is elevated in the cerebral cortex of *Ezh2* KO embryos^48^. Since the binding sites of PRC2 components to the genome in IPCs were not altered in *Auts2* cKO (Figure 8E), it seems unlikely that AUTS2 is involved in the genomic positioning of PRC2. AUTS2 might be involved in the H3K27 trimethylation activity of PRC2, but unfortunately, we have no direct evidence. We need to conduct additional analysis to clarify this issue.

In this study, we generated a mouse model for AUTS2 syndrome that exhibits a microcephaly-like phenotype. By analyzing model mice, we present pathomechanisms for microcephaly in AUTS2 syndrome and the molecular machinery to regulate IPC division and differentiation. We further suggest the new function of AUTS2 to suppress gene expression via H3K27 trimethylation. This study leads to a better understanding of the pathogenesis of AUTS2 syndrome and the regulatory mechanisms of gene expression and also provides insight into the evolution of the mammalian brain.

## Materials and Methods

### Animals

Generation of *Auts2^del8^* and *Emx1^Cre/+^; Auts2^flox^* alleles have been described previously^10,27^. *Auts2^del8/del8^* homozygotes are lethal at birth; therefore, the *Auts2^del8^* allele was maintained in heterozygosity in the C57BL6/N background. *Auts2^flox/flox^* and *Emx1^Cre/+^; Auts2^flox/flox^* mice were maintained on a mixed Balb/c and C57BL/6N background. Littermates of experimental mutants were used as controls. Pregnant Institute of Cancer Research (ICR) mice were purchased from Japan SLC Inc. (Shizuoka, Japan) and used for *in utero* electroporation. The day of the vaginal plug was considered embryonic day (E) 0.5. The mice were reared in a temperature-controlled, pathogen-free facility under a 12-h light/dark cycle with free access to food and water in ventilated racks. The Animal Care and Use Committee of the National Center for Neurology and Psychiatry approved all the animal experiments.

### Plasmids construction

The construction of pCAG-Myc-*Auts2*-full length (FL-*Auts2*), FL-*Auts2^R^* (shRNA-resistant FL-*Auts2*), NLS-FL-*Auts2^R^*, NES-FL-*Auts2^R^*, S-*Auts2*-var.1, S-*Auts2*-var.2 expression plasmids, pmU6-*Auts2,* and scrambled shRNA vectors have been described previously^27^. A cDNA fragment encoding full-length *Robo1* was amplified using a cDNA library from the E14.5 mouse brain. The fragment was inserted into a pCAGGS vector (GE Healthcare, Chicago, IL). shRNA vectors were generated by inserting double-stranded oligonucleotides into the mU6 pro vector. Target sequences are as follows; sh*Robo1* #1 (5′-GGATGATAAAGATGAAAGAAT -3′), sh*Robo1* #2 (5′-GGAAGTTACTGATGTGATTGC -3′), sh*Ezh2* (5′-GAAGTAAAGACTATGTTTAGT -3′), sh*Suz12* (5′-GAATTTAATGGAATGATTAAT -3′). Targeting sequences were designed using siDirect 2.0^49^. The shRNA-resistant *Ezh2* and *Suz12* expression vectors were constructed by introducing six silent mutations into cDNA sequence by PCR-mediated site-directed mutagenesis using the following primer set: sh*Ezh2*-resistant forward (5′-<u>G</u>GT<u>T</u>AA<u>A</u>AC<u>A</u>ATGTTGAGCTCCAATCGTCAGAAAATT -3′), sh*Ezh2*-resistant reverse (5′-<u>G</u>CT<u>C</u>AACAT<u>T</u>GT<u>T</u>TT<u>A</u>AC<u>C</u>TCATCAGCTCTTCTGAAC -3′), sh*Suz12*-resistant forward (5′-<u>G</u>TT<u>C</u>AA<u>C</u>GG<u>T</u>ATGAT<u>A</u>AA<u>C</u>GGAGAAACCAATGAAAAT -3′), sh*Suz12*-resistant reverse (5′-<u>G</u>TT<u>T</u>ATCAT<u>A</u>CC<u>G</u>TT<u>G</u>AA<u>C</u>TCTCTTCTTCCTGGACGA -3′) (Underlines indicate the mismatched nucleotide with shRNA). The pCAG-H3GFP vector was a gift from N. Masuyama. Plasmids were purified using an Endo-Free Plasmid Purification Kit (QIAGEN, Hilden, Germany).

### Immunofluorescence

Postnatal brains were dissected after perfusion fixation with 4% paraformaldehyde (PFA) in 0.1 M sodium phosphate buffer (pH 7.2) under deep isoflurane anesthesia. The brains were further fixed in 4% PFA for 2 h at room temperature (RT; 25±2 °C). Embryonic brains were fixed in 4% PFA for 3–4 h at 4 °C. Fixed brains were cryoprotected by immersion in 30% sucrose in PBS, embedded in optimum cutting temperature (Tissue-Tek O.C.T. compound, Sakura Fine-Tek, Tokyo, Japan) and cryosectioned at 14–16 µm (CM3050 S; Leica, Wetzlar, Germany). The points of the sections observed were set as follows: “rostral” is the anterior side where the corpus callosum is located, “central” is the anterior side where the hippocampus is located, and “caudal” is the posterior side where the hippocampus is located. Cells were fixed with 4% PFA for 20 min for immunocytochemistry at RT. Sections or cells were pre-incubated in blocking buffer containing 2–10% normal donkey serum (Merck Millipore, Burlington, MA) and 0.1% Triton X-100 at RT for 1 h and incubated overnight at 4 °C with primary antibodies and subsequently incubated with secondary antibodies conjugated with Alexa Fluor 488, Alexa Fluor 568, Alexa Fluor 594, or Alexa Fluor 647 (1:1000; abcam, Cambridge, UK or Jackson ImmunoResearch, West Grove, PA) and DAPI (5 µg/µl; Invitrogen, Carlsbad, CA) in blocking buffer for 1–2 h at RT. The following primary antibodies were used, rabbit anti-CUX1 (1:1000; sc-13024; Santa Cruz Biotechnology, Danvers, MA), rat anti-CTIP2 (1:3000; ab18465; abcam), rabbit anti-cleaved caspase-3 (1:400; 9661S; Cell Signaling Technology, Danvers, MA), rabbit anti-PAX6 (1:500; 901301; BioLegend, San Diego, CA), rabbit anti-TBR2 (1:1000; ab183991; abcam), rat anti-TBR2 (1:500; 14-4876-80; Invitrogen, Carlsbad, CA), rat anti-PH3 (pSer28) (1:400; H9908; Sigma-Aldrich, St. Louis, MO), rat anti-KI67 (1:500; 14-5698-82; Invitrogen), sheep anti-BrdU (1:200; ab1893; abcam), chicken anti-GFP (1:1000; ab13970; abcam), and goat anti-SOX2 (1:200; AF2018; R&D systems, Minneapolis, MN). For BrdU staining, before incubation with the blocking solution, the sections were incubated with 2 N HCl at 37 °C for 30 min. Fluorescent images were acquired using a laser scanning confocal microscope FV1000 (Olympus, Tokyo, Japan), SpinSR10 (Olympus), and a Zeiss LSM 780 confocal microscope system (Carl Zeiss, Oberkochen, Germany). The number of cells was measured using “Cell Counter” in ImageJ software.

### Nissl staining

Sections were stained with 0.1% cresyl violet in 1% acetic acid for 5 min, dehydrated using ethanol series and xylene, and mounted in Entellan. Stained sections were observed under a Keyence All-in-One microscope (BZ-X700, Osaka, Japan).

### EdU staining

Pregnant mice were intraperitoneally injected with 5-ethynyl-2’-deoxyuridine (EdU) (30 mg/kg) on E12.5, E15.5, and E16.5. Sections were stained using the Click-iT EdU imaging kit according to the manufacturer’s instructions (Thermo Fisher Scientific, Waltham, MA, USA).

### *In utero* electroporation

*In utero* electroporation experiments were performed as described previously with some modifications^50^. WT ICR, *Auts2^flox/flox^*, or homozygous *Auts2* cKO mice were anesthetized with isoflurane. pmU6-*Auts2*, *Robo1*, *Ezh2*, *Suz12* or scramble shRNA vector, pCAG-*Robo1* expression vector and pCAG-H3GFP vector were diluted to 1.5 µg/µl, 2 µg/µl and 0.5 µg/µl, respectively. For rescue experiments in Figure 3, indicated expression vectors (0.5 µg/µl) were coelectroporated with pmU6-*Auts2* shRNA vector and pCAG-H3EGFP vector. 1.5 µg/µl pCAG-*Ezh2*^R^ or *Suz12*^R^ vector was coelectroporated with pmU6-*Ezh2* or *Suz12* shRNA vector and pCAG-H3GFP vector. The plasmid solution with Fast Green (Sigma-Aldrich) was injected into the lateral ventricle of the mouse embryos and electroporated (37–40V, 50 ms, 450 ms intervals, five pulses) using a NEPA Gene Electroporator (NEPA21, Nepa Gene, Chiba, Japan).

### Calculation of cell cycle length

Based on previous methods^32,51^, the cell cycle length was calculated using BrdU and EdU. BrdU (50 mg/kg) and EdU (30 mg/kg) were administered intraperitoneally to pregnant mice 2 h and 30 min before sacrifice, respectively. The lengths of the S-phase (T_s_) and total cell cycle (T_c_) were calculated as follows: T_s_=1.5×S_cells_/L_cells_ and Tc=T_s_×P_cells_/S_cells_ [L_cells_=cells leaving S-phase (identified as TBR2^+^, BrdU^+^, EdU^−^ cells); S_cells_=cells in S-phase (TBR2^+^, EdU^+^ cells); P_cells_= total proliferating cells (TBR2^+^ cells)].

### Cell culture and transfection

HEK293T or Neuro2a cells were maintained in Dulbecco’s modified Eagle’s medium (DMEM, Sigma-Aldrich) supplemented with 10% fetal bovine serum (FBS) in a humidified atmosphere containing 5% CO_2_ at 37 °C. Plasmids were transfected into cells using Lipofectamine LTX Reagent (Invitrogen) following the manufacturer’s instructions. Transfected cells were cultured for 36–48 h.

### Western blotting analysis

Lysates of HEK293T cells and cerebral cortices from the mouse brain at E14.5 were solubilized in sodium dodecyl sulfate (SDS) sample buffer and separated by SDS-polyacrylamide gel electrophoresis. Proteins transferred onto a nitrocellulose membrane were immunoblotted with anti-TBR2 (1:1000; ab183991; abcam), ROBO1 (1:2000; 20219-1-AP; Proteintech, Rosemont, IL), and β-actin (1:500; M177-3; MBL, Nagoya, Japan) antibodies, and visualized using HRP-conjugated secondary antibodies (Jackson ImmunoResearch) followed by ECL Prime (GE Healthcare, Chicago, IL). Signals were detected using FUSION SOLO S (Vilber, Paris, France).

### Generation of *Tbr2^T2A-d2EGFP^* mice by CRISPR/Cas9

The method of KI mouse generation has been described previously^52^. The coding sequences of the inserted T2A (shown in lower case) and d2EGFP (shown in upper case) before the stop codon of *Tbr2* locus are as follows: 5′-ggcagtggagagggcagaggaagtctgctaacatgcggtgacgtcgaggagaatcctggcccaATGGTGAGCAAGGGCGAGGAGCTGTTCACCGGGGTGGTGCCCATCCTGGTCGAGCTGGACGGCGACGTAAACGGCCACAAGTTCAGCGTGTCCGGCGAGGGCGAGGGCGATGCCACCTACGGCAAGCTGACCCTGAAGTTCATCTGCACCACCGGCAAGCTGCCCGTGCCCTGGCCCACCCTCGTGACCACCCTGACCTACGGCGTGCAGTGCTTCAGCCGCTACCCCGACCACATGAAGCAGCACGACTTCTTCAAGTCCGCCATGCCCGAAGGCTACGTCCAGGAGCGCACCATCTTCTTCAAGGACGACGGCAACTACAAGACCCGCGCCGAGGTGAAGTTCGAGGGCGACACCCTGGTGAACCGCATCGAGCTGAAGGGCATCGACTTCAAGGAGGACGGCAACATCCTGGGGCACAAGCTGGAGTACAACTACAACAGCCACAACGTCTATATCATGGCCGACAAGCAGAAGAACGGCATCAAGGTGAACTTCAAGATCCGCCACAACATCGAGGACGGCAGCGTGCAGCTCGCCGACCACTACCAGCAGAACACCCCCATCGGCGACGGCCCCGTGCTGCTGCCCGACAACCACTACCTGAGCACCCAGTCCGCCCTGAGCAAAGACCCCAACGAGAAGCGCGATCACATGGTCCTGCTGGAGTTCGTGACCGCCGCCGGGATCACTCTCGGCATGGACGAGCTGTACAAGAAGCTTAGCCATGGCTTCCCGCCGGAGGTGGAGGAGCAGGATGATGGCACGCTGCCCATGTCTTGTGCCCAGGAGAGCGGGATGGACCGTCACCCTGCAGCCTGTGCTTCTGCTAGGATCAATGTGTAG-3′. The left (5′-GGAAACTCGCCCCCCATAAAGTGTGAGGACATTAACACTGAAGAGTACAGTAAAGACACCTCCAAAGGCATGGGGGCTTATTATGCTTTTTACACAAGTCCC -3′) and right (5′-GGATACATCAAAGGTGGAAGGCAAAAGTCTTTTTGGTAACCTAGGCAAAGAACACAACAAAACACCACCAGGTCCATCTGGAAAGGTTAAAGGTTAAAATAAT GCT -3′) homology arms were connected to the insertion sequence.

Cas9 protein, crRNA/tracrRNA, and donor ssDNA were microinjected into B6C3F1 mouse zygotes following standard protocols. KI mice were screened using PCR with a pair of primers and sequence analyses, forward primer (5′-TAACGGTGAGAGAACCGTGC-3′) and reverse primer (5′-AAAACACTCCTGCGTCCTCC-3′). The founders were backcrossed with C57BL/6N mice. The following three primers were used for genotyping the progenies using GoTaq DNA Polymerase (Promega, Madison, WI): F; 5′-GGTGTACAACAGCGCTTGCA-3, R1; 5′-GACGTCACCGCATGTTAGCA-3′, R2; 5′-TTTGGCGCCTTCTCTCAGAG-3′.

### *In vitro* digestion assay

The genomic region (813 bp) containing the *Tbr2*-crRNA target sequences was PCR amplified using PrimeSTAR Max (Takara Bio, Shiga, Japan), and a pair of primers, F (5ʹ- TAACGGTGAGAGAACCGTGC-3ʹ) and R (5ʹ-AAAACACTCCTGCGTCCTCC-3ʹ). Cas9 protein (50 ng/µL), *Tbr2*-crRNA (15.8 ng/µL; 5′-AAAGGUUAAAAUAAUGCUCUAGGGUUUUAGAGGUAUGCUCUUUUG-3′), and tracrRNA (25.8 ng/µL; 5′-AAACAGCAUAGCAAGUUAAAAUAAGGCUAGUCCGUUAUCAACUUGAAAAAGUGGCACCGAGUCGGUGCU-3′) were incubated with the *Tbr2* target PCR products in Cas9 Nuclease Reaction Buffer for 60 min at 37°C. Reactions were stopped using 10X DNA loading buffer containing 40% glycerol, 2% SDS, and 180 mM EDTA. The samples were analyzed by electrophoresis using 1.5% agarose gels.

### Cell sorting

The lateral parts of the cerebral cortices of the embryos were dissected and dissociated using a neuron dissociation solution (Fujifilm Wako Pure Chemical Corporation, Osaka, Japan). Dissociated cells were stained with Allophycocyanin (APC)-conjugated antibodies against CD133 (1:400; 141207; BioLegend). Cells were sorted on a FACS Aria fusion cell sorter (Becton Dickinson, Tokyo, Japan) with a 100-µm nozzle.

### RT-qPCR

Total RNA was extracted using an RNeasy Micro Kit (QIAGEN). Purified total RNA was reverse-transcribed into cDNA using the ReverTra Ace qPCR RT kit (Toyobo, Osaka, Japan) according to the manufacturer’s instructions. Real-time qPCR was performed with PowerUp SYBR Green Master Mix (Thermo Fisher Scientific) in a Light Cycler 96 (Roche Diagnostics, Basel, Switzerland), and relative expression levels were calculated using the 2Δ method. The quantified amount of target mRNA was normalized to that of internal control *Actb* mRNA. Primers used are listed in Supplementary Table 3.

### Co-immunoprecipitation

Nuclei were extracted with Nuclear Extraction Buffer (20 mM HEPES-KOH pH 7.9, 10 mM KCl, 0.1% Triton X-100, 20% Glycerol, 0.5 mM Spermidine, 1x Protease Inhibitor Cocktail (Roche)) from transfected Neuro2a cells or E14.5 cerebral cortices. The nuclei were dissolved with lysis buffer (50 mM Tris-HCl, pH7.5, 125 mM NaCl, 1 mM EDTA, 5% glycerol, 0.5% NP-40, 1 mM PMSF and 1x Protease Inhibitor Cocktail (Roche), 100 µg/µl RNaseA). The lysate was sonicated using a Bioruptor (Five cycles of 30 s ON and 30 s OFF) (SonicBio, Kanagawa, Japan). After centrifugation, the supernatant was incubated with mouse anti-FLAG (F1804; Sigma-Aldrich), rabbit anti-AUTS2 (HPA00390; Sigma-Aldrich) or rabbit IgG (011-000-003; Jackson ImmumoResearch)-conjugated dynabeads protein G (Thermo Fisher Scientific) for 4 h at 4 °C. After washing with lysis buffer, all precipitates were solubilized in SDS sample buffer and detected by western blotting with rabbit anti-FLAG (1:1000; F7425; Sigma-Aldrich), rabbit anti-EZH2 (1:1000; 5246S; Cell Signaling Technology), rabbit anti-SUZ12 (1:1000; 3737S; Cell Signaling Technology) and rabbit anti-AUTS2 (1:500; HPA00390; Sigma-Aldrich) antibodies.

### Proximity ligation assay

EGFP^+^ cells were plated on cover grasses coated with 1 mg/ml poly-D-lysine (Sigma-Aldrich) for 1 h in a humidified atmosphere containing 5% CO_2_ at 37 °C. Cells were fixed with 4% PFA for 20 min and permeabilized with 0.2% Triton X-100 for 15 min at RT. The Duolink Proximity Ligation Assay (Sigma-Aldrich) was performed following the manufacturer’s instructions. Cells were incubated overnight at 4 °C with primary antibodies as follows; rabbit anti-AUTS2 antibody (1:400; HPA00390; Sigma-Aldrich) and mouse anti-EZH2 antibody (1:100; 3147S; Cell Signaling Technology). Signals were acquired using a super-resolution confocal microscope SpinSR10 (Olympus).

### RNA-seq

Total RNA was extracted from sorted cells from six control (*Auts2^flox/flox^*) and six *Auts2* cKO (*Emx1^Cre/+^; Auts2^flox/flox^*) mice at E15.5, using an RNeasy Micro Kit (QIAGEN). Quality analyses and quantification of the extracted RNA were performed using a NanoDrop and Qubit Fluorometer (Thermo Fisher Scientific). Sequencing libraries were prepared using the NEBNext Ultra II Directional RNA Library Prep Kit (New England BioLabs, Ipswich, MA) according to the manufacturer’s instructions. RNA-seq libraries were sequenced using the Illumina HiSeq 4000 platform.

The obtained reads were processed using fastp to trim adaptor sequences and remove low-quality nucleotides at the read ends^53^. The processed reads were aligned to the reference mouse genome (GRCm38/mm10) using HISAT2^54^. Genome-wide expression levels were measured as transcripts per kilobase million (TPM) using StringTie^55^. The number of reads per gene was counted for each sample using StringTie^55^. Differentially expressed genes (DEGs) were identified using DESeq2^56^. Gene Ontology biological process analysis was performed using the DAVID bioinformatics Resources (National Institute of Allergy and Infectious Diseases, National Institute of Health; https://david.ncifcrf.gov).

### CUT&Tag

CUT&Tag was performed as previously described with some modifications^36^. Briefly, 100,000 EGFP^+^ cells sorted by FACS were used for each experimental replicate. Cell nuclei were extracted with Nuclear Extraction Buffer (20 mM HEPES-KOH pH 7.9, 10 mM KCl, 0.1% Triton X-100, 20% Glycerol, 0.5 mM Spermidine, 1x Protease Inhibitor Cocktail (Roche)), bound to wheat germ agglutinin (WGA) coated magnetic beads (Bangs Laboratories, Fishers, IN), and incubated for overnight at 4 °C with a primary antibody in Antibody Binding Buffer (20 mM HEPES pH 7.5, 150 mM NaCl, 0.5 mM Spermidine, 2 mM EDTA, 0.01% Digitonin, 1x Protease Inhibitor Cocktail (Roche)). The following primary antibodies were used, rabbit anti-AUTS2 (HPA000390; Sigma-Aldrich), rabbit anti-RING1B (5694S; Cell Signaling Technology), rabbit anti-H2AK119ub (8240S; Cell Signaling Technology), rabbit anti-H3K27me3 (9733S; Cell Signaling Technology), rabbit anti-H3K4me3 (9751S; Cell Signaling Technology), rabbit anti-H3K27ac (8173S; Cell Signaling Technology), rabbit anti-EZH2 (5246S; Cell Signaling Technology), and rabbit anti-SUZ12 (3737S; Cell Signaling Technology). The beads were incubated with a secondary antibody (EpiCypher, Durham, NC) in a Dig-wash Buffer (20 mM HEPES pH 7.5, 150 mM NaCl, 0.5 mM Spermidine, 0.01% Digitonin, 1x Protease Inhibitor Cocktail (Roche)) for 30 min at RT. The beads were washed twice in Dig-wash Buffer and incubated with protein A/B–Tn5 conjugates (EpiCypher) in Dig-wash 300 Buffer (20 mM HEPES pH 7.5, 300 mM NaCl, 0.5 mM Spermidine, 0.01% Digitonin, 1x Protease Inhibitor Cocktail) for 1 h at RT. After washing twice with Dig-wash 300 Buffer, the beads were resuspended with Tagmentation Buffer (20 mM HEPES pH 7.5, 300 mM NaCl, 0.5 mM Spermidine, 0.01% Digitonin, 10 mM MgCl_2_, 1x Protease Inhibitor Cocktail (Roche)) and incubated for 1 h at 37 °C. Furthermore, 0.5 M EDTA (final 17 mM), 10% SDS (final 0.1%), and 20 mg/ml Proteinase K (final 0.17 mg/ml) were added to 100 µl of the sample and incubated at 55 °C for 1 h to stop tagmentation. Sequencing libraries were amplified using NEBNext HiFi 2x Master Mix (NEB) as follows: 72 °C for 5 min and 98 °C for 30 s followed by 15 cycles (for histone modifications), 18 cycles (for AUTS2 and RING1B) or 20 cycles (for EZH2 and SUZ12) of 10 s at 98 °C and 10 s at 63 °C. The DNA fragments and libraries were purified using a DNA cleaner and concentrator (ZYMO Research, Irvine, CA). For input, EGFP^+^ cells were sonicated with a Bioruptor (25 cycles of 30 s ON and 30 s OFF) (SonicBio), and the DNA fragment was purified. Input libraries were prepared using ThruPLEX (Takara Bio) following the manufacturer’s instructions. The size distribution and concentration of libraries were determined using Agilent 4150 TapeStation analysis (Agilent Technologies, Santa Clara, CA). Deep sequencing of CUT&Tag was performed using 150-bp paired-end sequencing on the Illumina HiSeq X platform.

### CUT&Tag data analysis

Paired-end reads were processed with fastp to trim adaptors and remove short- and low-quality reads^53^, followed by alignment to the reference mouse genome mm10 using Bowtie2^57^. SAM files were converted to the BAM format using SAMtools^58^. PCR duplicates were removed using Picard Mark Duplicates (https://github.com/broadinstitute/picard). Peaks were identified using an IDR pipeline (https://hbctraining.github.io/Intro-to-ChIPseq/lessons/07_handling-replicates-idr.html) and MACS2^59^ (parameters: for AUTS2 and SUZ12; -q 0.05, for the others; –q 0.01) from pseudo-replicates with the input library as a control. Genomic peak annotation was performed using the R package ChIPseeker^60^. Genomic annotation was performed using GREAT software^61^. Reads were normalized to one million paired-end reads for histone modifications or 100 thousand paired-end reads for AUTS2, RING1B, EZH2, and SUZ12 to generate BigWig files using bamCompare from deepTools^62^. Heat maps and density plots were generated using DeepTools. Log_2_ enrichment of CUT&Tag signals was performed using multiBigwigSummary in BED file mode from deepTools. Bed files of the TSS of upregulated or unchanged genes were generated using UCSC (https://genome.ucsc.edu/). BigWig files were visualized using the Integrative Genomics Viewer (IGV) ^63^. We merged three independent samples into one sample with the cat command, followed by the above analysis using three merged samples to supplement the read coverage of the EZH2 and SUZ12 libraries.

### Processing of scRNA-seq data

Count datasets of only cortical cells at E13.5 and E15.5 were downloaded from GSE107122 and processed with the Seurat R package as described^64^. For clustering, the parameter “resolution = 0.5” was used. Annotation of each cluster was performed using representative marker genes such as *Sox2*, *Eomes*, *Neurod1*, *Tbr1*, *Bcl11b*, *Satb2,* and *Pou3f2*.

### Statistical analysis

Statistical analyses, with the exception of RNA-seq, were performed using GraphPad Prism 7 software (GraphPad Software, La Jolla, CA, USA). No statistical methods were used to pre-determine the sample size; however, the sample size was determined based on previous studies^27,32,33,65,66,67^. Sample size was indicated in the figure legends. The normal distribution of data was confirmed using the Shapiro-Wilks test. If significant, the data was compared by a nonparametric Mann-Whitney U test. The data with normal distribution were compared by two-tailed unpaired t-test. For comparison more than 2 groups, the normally distributed data was used a one-way analysis of variance (ANOVA) followed by the Dunnett’s or Turkey’s multiple comparison test, and the non-normally distributed data was used a Kruskal–Wallis test. No randomization of samples was performed. Data analyses were not performed blinded to the genotype. Statistical significance was set at a P-value < 0.05.

## Data and Materials Availability

RNA-seq data and CUT&Tag data have been deposited into the DNA Data Bank of Japan (DDBJ) database with the accession number DRA017450 and DRA017448, respectively. Source data are provided with this paper. KI mouse line generated in this study and used research materials are available on request to M.H.

## Code availability

Every code used in this study was executed with default settings and the parameters indicated in the Materials and Methods sections. Detailed codes used in this study are available upon request to M.H.

## Acknowledgements

We thank Dr. Kishi (The University of Tokyo) for the guidance on the use of FACS. This work was supported by Grants-in-Aid for Scientific Research, KAKENHI (Grants 16H06531 to Y.G., 22K15134 to K.S. and 22H02730 to M.H.); AMED (Grant JP23wm0425005h0003 to M.H.), an Intramural Research Grant of NCNP (3-9, 4-5, 4-6 to M.H.); Japan Health Research Promotion Bureau (JH) under Research Fund (2020-B-07 to M.H.); and Tokumori Yasumoto Memorial Trust (M.H.).

## Author contributions

K.H. and M.H supervised this project. K.Shimaoka. designed the study. K.Shimaoka., K.H., and M.H. wrote the manuscript. K.Shimaoka., A.S., I.H., K.Y., and S.F.E. performed imaging experiments and statistical analysis. S.T., Y.G., K.Shimaoka., and S.M. performed RNA-seq and analyzed the data. K.Shimaoka. and T.Imamura. performed and supervised CUT&Tag experiments and the data analysis. Y.U.I., and T.Inoue. generated and supervised the designs of *Eomes^T2A-d2EGFP^* KI mice. M.A. and K.Sakimura. generated and supervised the designs of *Auts2* mutant mice.

## Supplementary Information

Supplementary Table 1: DEGs identified by RNA-seq. Layer1–DEGs between CD133^high^ cells and EGFP^+^ cells from control cortices. Layer2–DEGs in EGFP^+^ cells from *Auts2* cKO cortices. Layer3–DEGs in CD133^high^ cells from *Auts2* cKO cortices.

Supplementary Table 2: List of significant GO terms for DEGs (P-value < 0.05). Layer1–GO terms for upregulated genes in EGFP^+^ cells. Layer2–GO terms for downregulated genes in EGFP^+^ cells. Layer3–GO terms for upregulated genes in CD133^high^ cells. Layer4–GO terms for downregulated genes in CD133^high^ cells.

Supplementary Table 3: List of primers.

---

**Figure S1:**
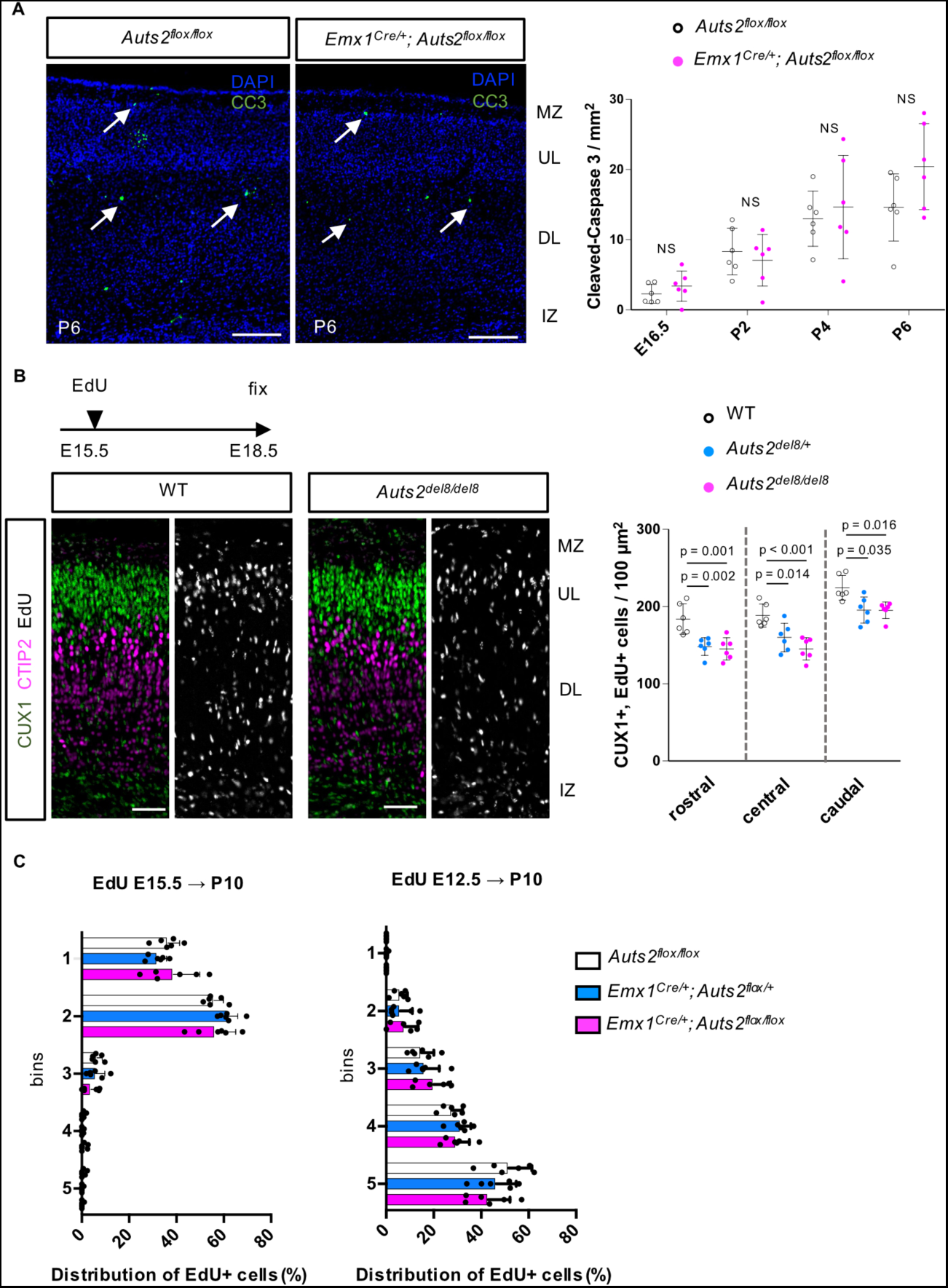
AUTS2 regulates the production of neurons but not apoptosis. (A) Immunostaining for cleaved-caspase3 (CC3, green) and DAPI (blue) on the cerebral cortical sections of *Auts2^flox/flox^*and *Emx1^Cre/+^; Auts2^flox/flox^* mice at P6. Arrows indicate CC3^+^ cells. The graph shows the number of CC3^+^ cells in the cerebral cortex in the *Auts2^flox/flox^* and *Emx1^Cre/+^; Auts2^flox/flox^* mice at E16.5, P2, P4, and P6. MZ, marginal zone; UL, upper-layer; DL, deep-layer; IZ, intermediate zone. (B) One pulse EdU labeling was performed in E15.5 WT, *Auts2^del8/+^*, and *Auts2^del8/del8^* mice. The embryos were analyzed at E18.5 by triple-staining with CUX1 (green), CTIP2 (magenta), and EdU (white). Representative images show the rostral sections of cerebral cortices in WT and *Auts2^del8/del8^* mice. The graph shows the number of CUX1/EdU-double positive cells in a 100 µm^2^ area of WT, *Auts2^del8/+^*, and *Auts2^del8/del8^* cerebral cortex. (C) Distribution of EdU^+^ cells labeled at E15.5 (left) and E12.5 (right) in *Auts2^flox/flox^*, *Emx1^Cre/+^; Auts2^flox/+^*and *Emx1^Cre/+^; Auts2^flox/flox^* cerebral cortices at P10. The graphs show the percentage of EdU^+^ cells at the rostral point in the five bins relative to total EdU^+^ cells, as indicated in Figures 1F and G. Bin 1 includes the marginal zone and upper layer, bin 2 is the upper layer, bin 3 is the upper and deep layers, and bins 4–5 are the deep layers. Data are presented as the mean ± SD (N=three mice, six sections). NS, not significant, Student’s t-test (A) and One-way ANOVA with Dunnett’s post-hoc test or Kruskal– Wallis test (B, C). Scale bars, 200 µm (A) and 100 µm (B).

**Figure S2:**
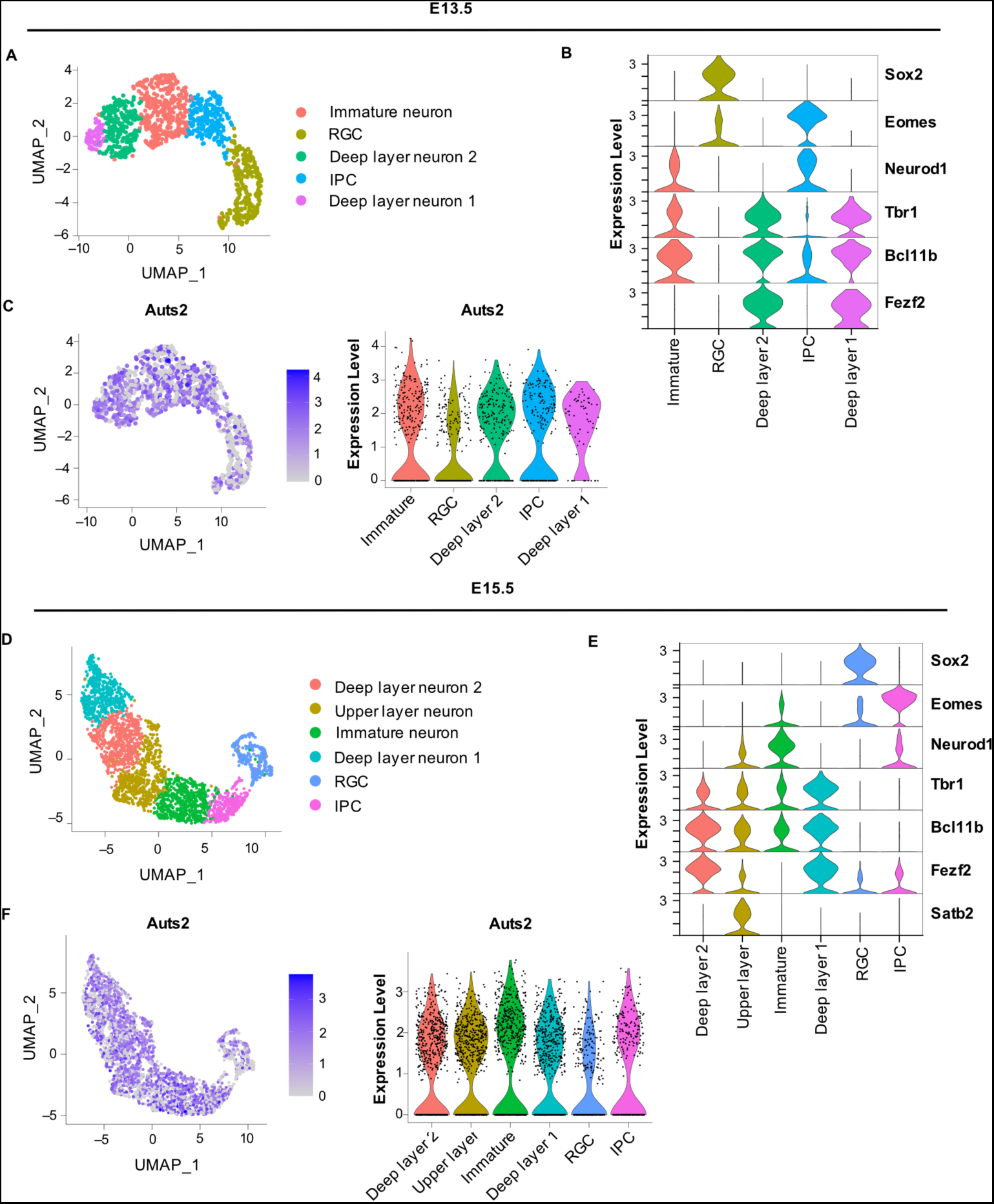
Expression of *Auts2* transcripts in developing mouse cortical cells. Data of WT cerebral cortical cells at E13.5 (A–C) and E15.5 (D–F) from previously published scRNA-seq data^30^ were processed with the computational pipeline (Seurat). (A, D) UMAP-based dimensional reduction and cluster analysis grouped five users for E13.5 (A) and six clusters for E15.5 (D). (A) (B) Violin plots showing the expression levels of representative marker genes (*Sox2*, *Eomes*, *Neurod1*, *Tbr1*, *Bcl11b,* and *Fezf2*). Each cluster was annotated based on these expressions (C, F) Expression levels of *Auts2* transcripts are shown in feature and violin plots at E13.5 (C) and E15.5 (F). (E) Violin plots showing the expression levels of representative marker genes (*Sox2*, *Eomes*, *Neurod1*, *Tbr1*, *Bcl11b*, *Fezf2,* and *Satb2*). Each cluster was annotated based on these expressions.

**Figure S3:**
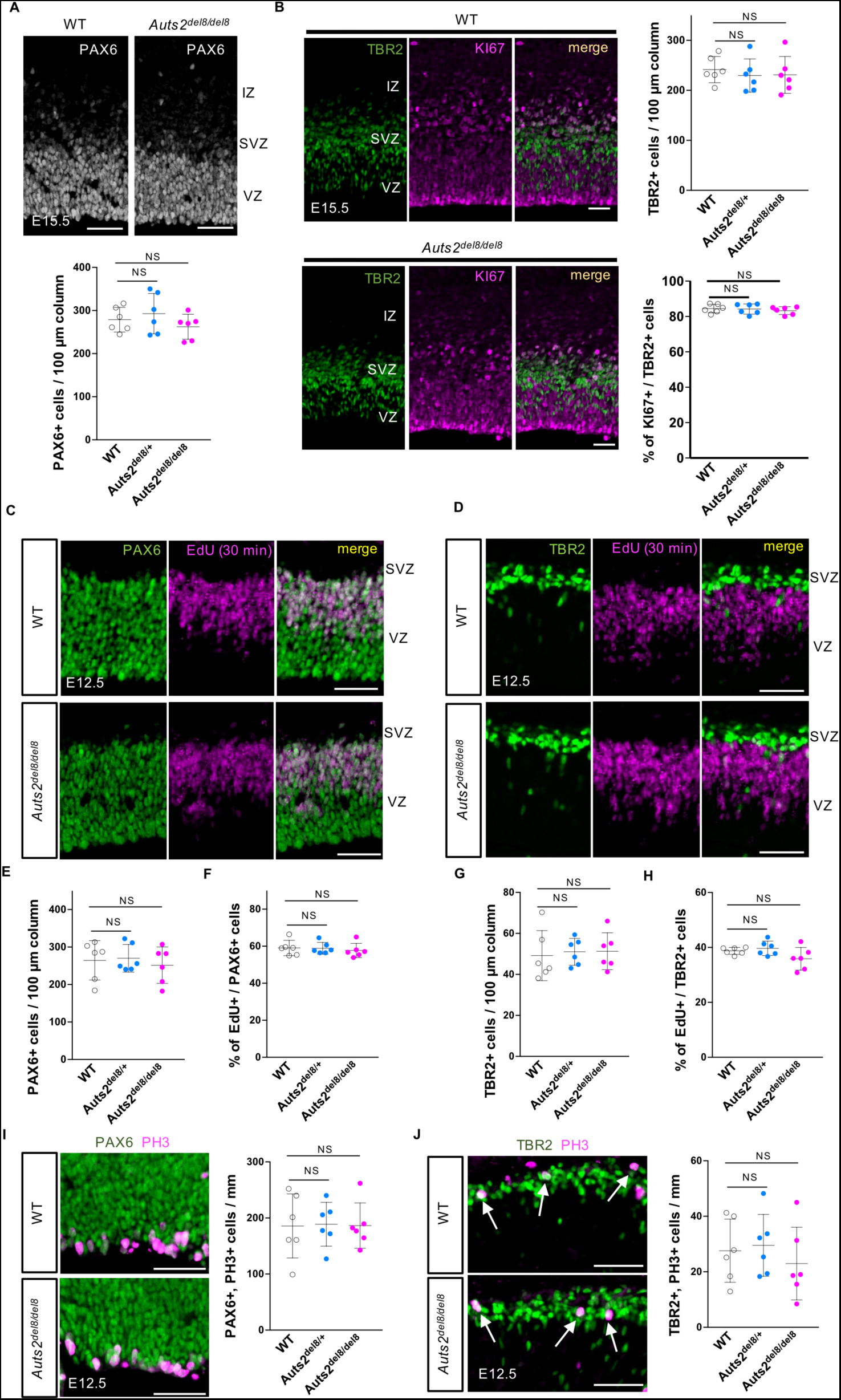
Loss of *Auts2* does not affect the proliferation of RGCs and IPCs at E12.5. (A) Representative images of immunostaining for PAX6 in WT and *Auts2^del8/del8^* cortical sections at E15.5. The graph shows the number of PAX6^+^ cells within a 100 µm wide field. (B) Representative images of immunostaining for TBR2 (green) and KI67 (magenta) in WT and *Auts2^del8/del8^*cortical sections at E15.5. The graph shows the number of TBR2^+^ cells within a 100 µm wide field (top) and the ratio of KI67^+^ cells in TBR2^+^ cells (bottom) (C–H) EdU was administered intraperitoneally to pregnant mice 30 min before sacrifice at E12.5. (D) Representative staining images for PAX6 (green) and EdU (magenta) in WT and *Auts2^del8/del8^* cortical sections. (E) Representative staining images for TBR2 (green) and EdU (magenta) in WT and *Auts2^del8/del8^*cortical sections. (E, G) The graphs show the number of PAX6^+^ (E) and TBR2^+^ (G) cells within a 100 µm wide field. (F, H) The graphs show the percentage of EdU^+^ cells in PAX6^+^ cells (F) and TBR2^+^ cells (H). (I) Representative Immunostaining images for PAX6 (green) and PH3 (magenta) in WT and *Auts2^del8/del8^* cortical sections at E12.5. The graph shows the number of PAX6^+^ and PH3^+^ cells on the ventricular surface in WT, *Auts2^del8/+^*, and *Auts2^del8/del8^*mice. (J) Representative Immunostaining images for TBR2 (green) and PH3 (magenta) in WT and *Auts2^del8/del8^* cortical sections at E12.5. The graph shows the number of TBR2^+^ and PH3^+^ cells in WT, *Auts2^del8/+^*, and *Auts2^del8/del8^* mice. Arrows indicate TBR2^+^ and PH3^+^ cells. The number of cells was quantified at the rostral point. Data are presented as the mean ± SD (N = three mice, six sections). NS, not significant, One-way ANOVA with Dunnett’s post-hoc test. Scale bars, 50 µm. VZ, ventricular zone; SVZ, subventricular zone; IZ, intermediate zone.

**Figure S4:**
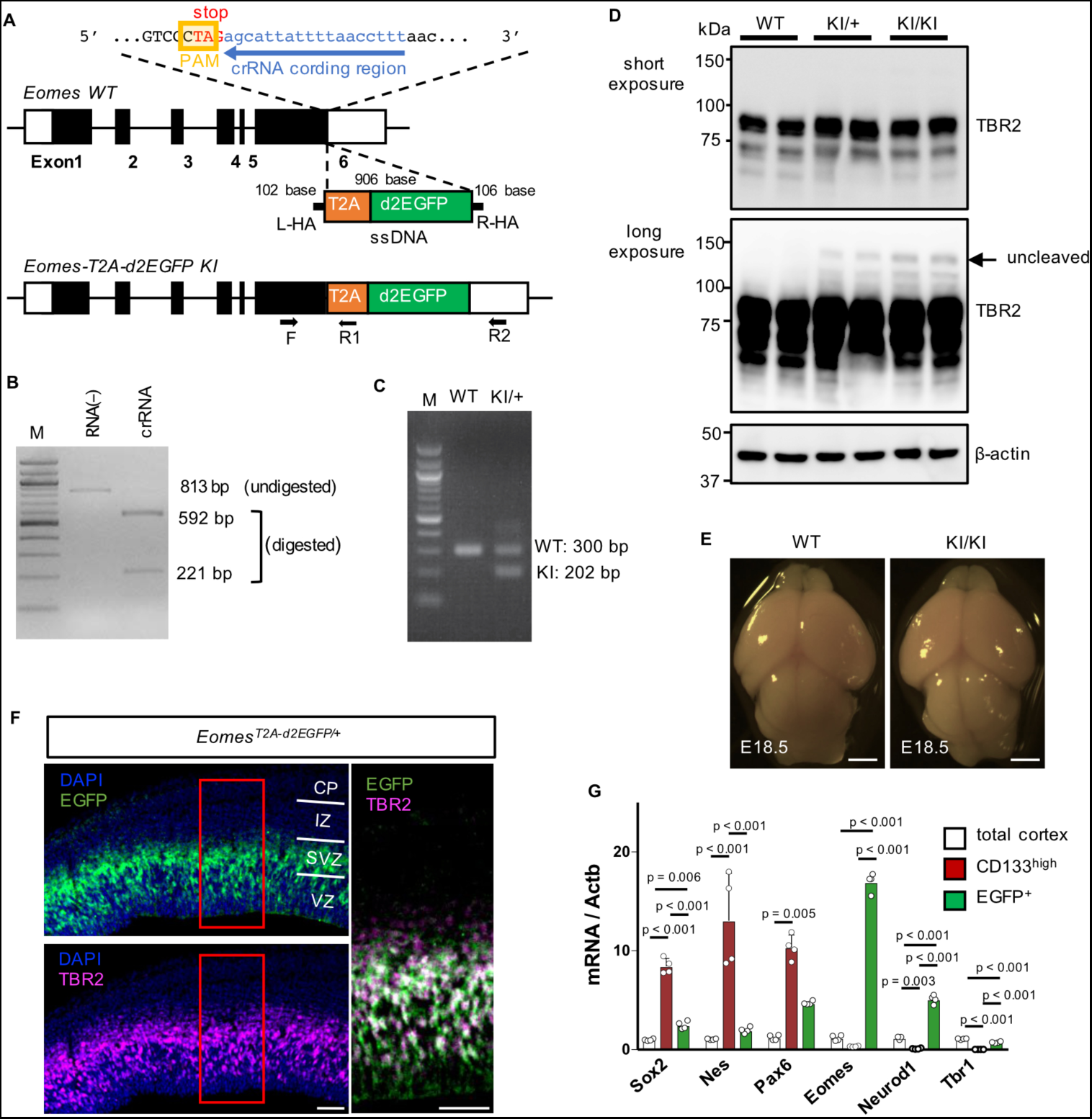
Generation of *Eomes^T2A-d2EGFP^*knock-in mouse. (A) Targeting strategy of CRISPR/Cas9 method for the generation of *Eomes^T2A-d2EGFP^* knock-in mouse. The DNA sequences for T2A-d2EGFP were inserted between the last amino acid and the stop codon of the *Eomes* locus. L-HA, left homology arm; R-HA, right homology arm; ssDNA, single-stranded donor DNA; F, forward primer for genotyping; R1 and R2, reverse primers for genotyping. (B) *In vitro* digestion assay to check the cleavage activity of the crRNA designed in (A). Targeted PCR products of the *Eomes* locus were cleaved in the presence of the chemically synthesized guide RNA (*Eomes* crRNA/tracrRNA) combined with Cas9 protein. M, molecular marker. (C) Genotyping for WT and *Eomes^T2A-d2EGFP/+^*(KI) mouse using primers indicated in (A). (D) Western blotting analysis of cerebral cortical lysates in WT, *Eomes^T2A-d2EGFP/+^*(KI/+), and *Eomes^T2A-d2EGFP/T2A-d2EGFP^* (KI/KI) mice at E14.5 with anti-TBR2 and anti-ß-actin antibodies. A very small amount of uncleaved TBR2-T2A-d2EGFP protein was observed at the long exposure (arrow); however, most signals for TBR2 were detected around the expected size for TBR2 protein (short exposure), indicating that TBR2 and d2EGFP proteins were efficiently cleaved in those mice. (E) Whole-mount images in WT and *Eomes^T2A-d2EGFP/T2A-d2EGFP^* (KI/KI) mouse brains at E18.5. Scale bars, 1 mm. (F) Representative images of staining with DAPI (blue), anti-GFP (green), and anti-TBR2 (magenta) antibodies in *Eomes^T2A-d2EGFP/+^* cortical sections at E14.5. CP, cortical plate; IZ, intermediate zone; SVZ, subventricular zone; VZ, ventricular zone. Scale bars, 50 µm. (G) RT-qPCR analysis for *Sox2, Nes, Pax6, Eomes, Neurod1* and *Tbr1* in the cerebral cortex, sorted CD133^high^ cells and sorted EGFP^+^ cells from *Eomes^T2A-d2EGFP/+^* mice at E15.5. Data are presented as mean ± SD (N = four biological replicates); One-way ANOVA with Turkey’s post-hoc test or Kruskal–Wallis test.

**Figure S5:**
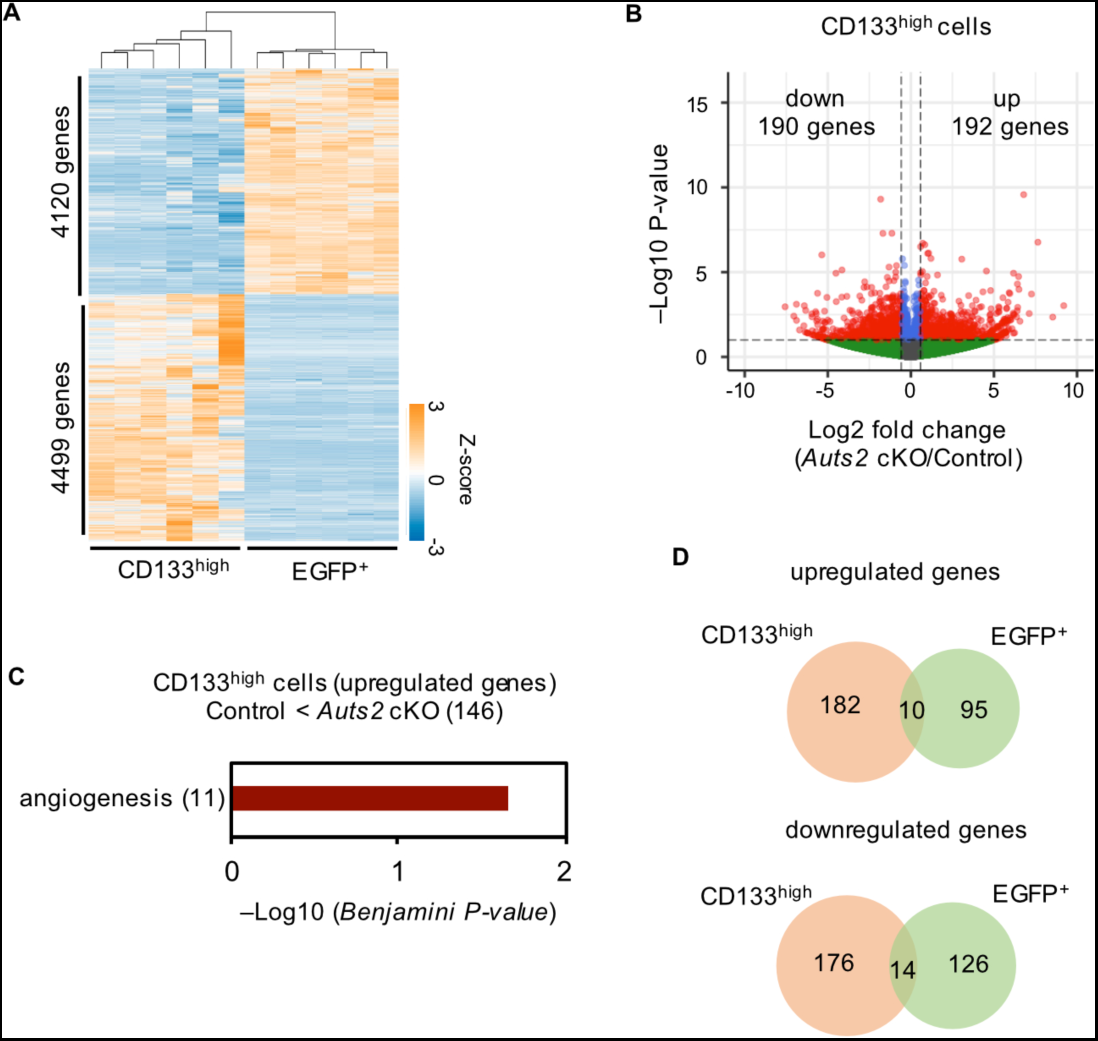
Transcriptional profiling of RGCs at E15.5. (A) Heatmap of differentially expressed genes (DEGs) between CD133^high^ and EGFP^+^ cells from E15.5 control mouse cortices (*Padj* <0.05). In total, 4499 and 4120 genes were enriched in CD133^high^ and EGFP^+^ cells, respectively. The color scale is shown on the bottom right. (B) Volcano plot showing differences in gene expression in CD133^high^ cells between control and *Auts2* cKO mice. Red plots indicate upregulated and downregulated genes in *Auts2* cKO cells compared with the control (P-value <0.01 and |Log2 fold-change| > 0.58). (C) DAVID Gene Ontology biological process analysis of upregulated genes in CD133^high^ cells (Benjamini-Hochberg adjusted P-value < 0.05). Numbers in parentheses show the count of genes. (D) Venn diagrams showing upregulated or downregulated genes overlap between CD133^high^ and EGFP^+^ cells.

**Figure S6:**
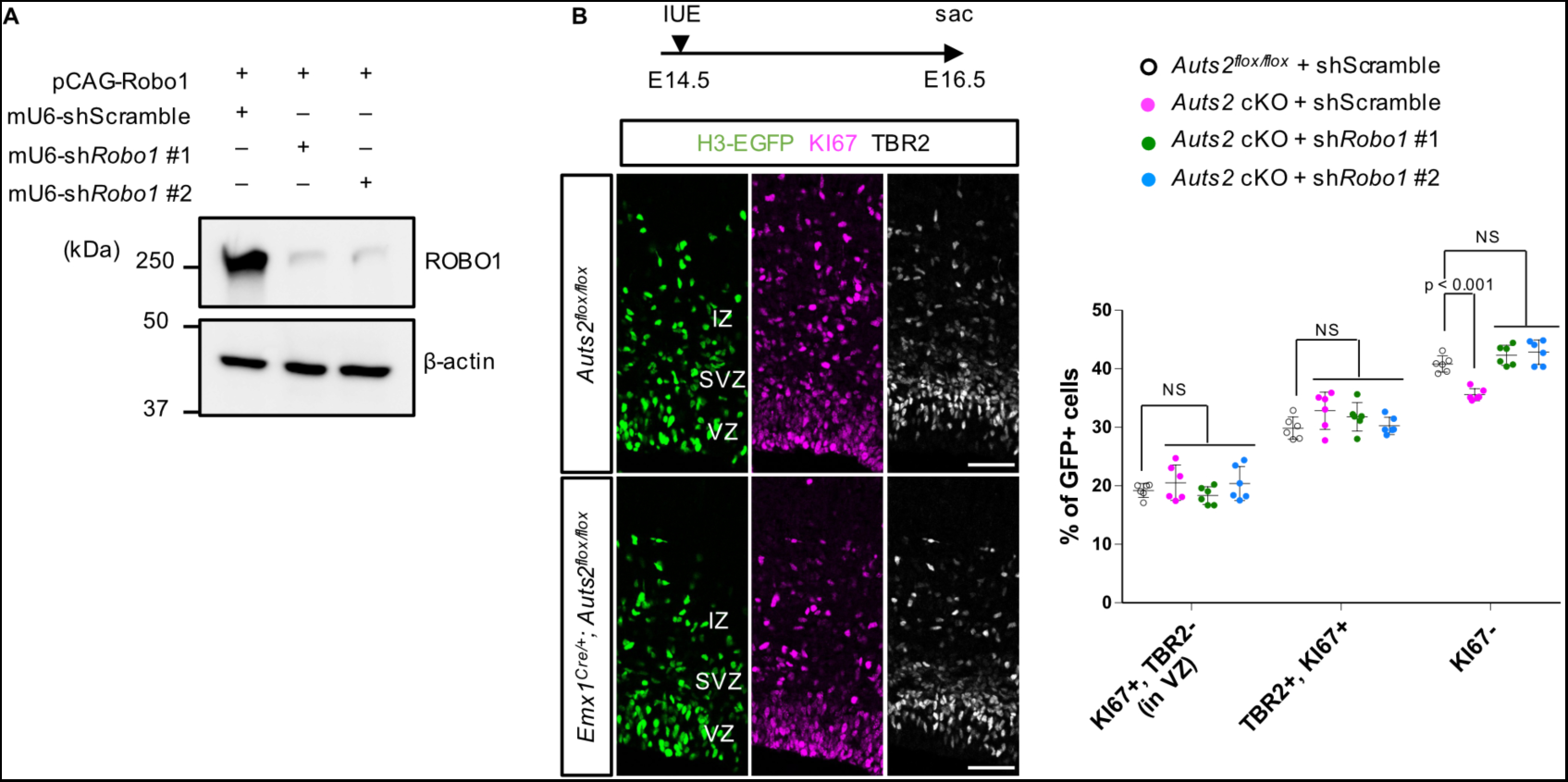
The efficiency of *Robo1* shRNAs and rescue experiments by *Robo1* knockdown. (A) *In vitro* knockdown (KD) experiment of ROBO1 expression by co-transfection of a *Robo1* expression vector and scrambled shRNA or *Robo1* shRNA vectors into HEK293 cells. The cell lysates were immunoblotted with anti-ROBO1 and anti-ϕ3-actin antibodies 2 days after transfection. (B) *In utero* electroporation (IUE) of scrambled or *Robo1* shRNA into *Auts2^flox/flox^* or *Emx1^Cre/+^; Auts2^flox/flox^* (*Auts2* cKO) cortices at E14.5, followed by immunostaining with anti-GFP (green), anti-KI67 (magenta) and anti-TBR2 (white) at E16.5. The graph shows the proportion of KI67^+^ and TBR2-negative cells located in the VZ (RGCs), TBR2^+^ and KI67^+^ cells (IPCs), and KI67-negative cells (postmitotic neurons) among total GFP^+^ cells. The percentage of RGCs and IPCs in electroporated cells was not different among the indicated samples. Data are presented as the mean ± SD (N = three mice, six sections). NS, not significant, One-way ANOVA with Dunnett’s post-hoc test. Scale bars, 50 µm.

**Figure S7:**
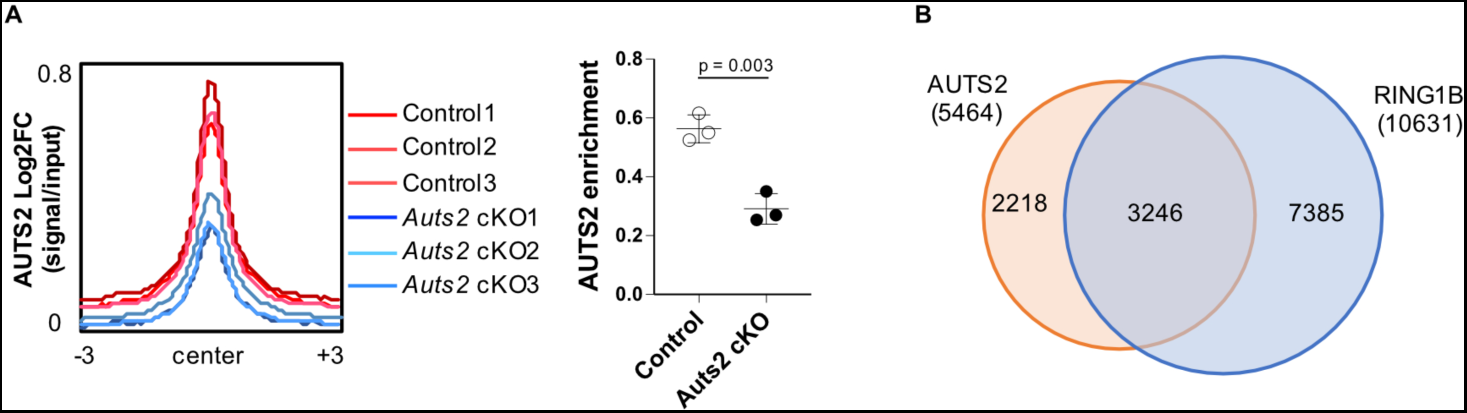
CUT&Tag signals of AUTS2 reduced in *Auts2* mutants. (A) Density plots of AUTS2 CUT&Tag signals in control and *Auts2* cKO homozygous cells centered on AUTS2-binding loci (±3 kb). The graph shows the enrichment of CUT&Tag signals for AUTS2 in control and *Auts2* cKO cells. FC, fold change. N = three biological replicates. Data are presented as the mean ± SD; unpaired Student’s t-test. (B) Venn diagrams comparing the AUTS2-binding loci with RING1B-binding loci in control cells.

**Figure S8:**
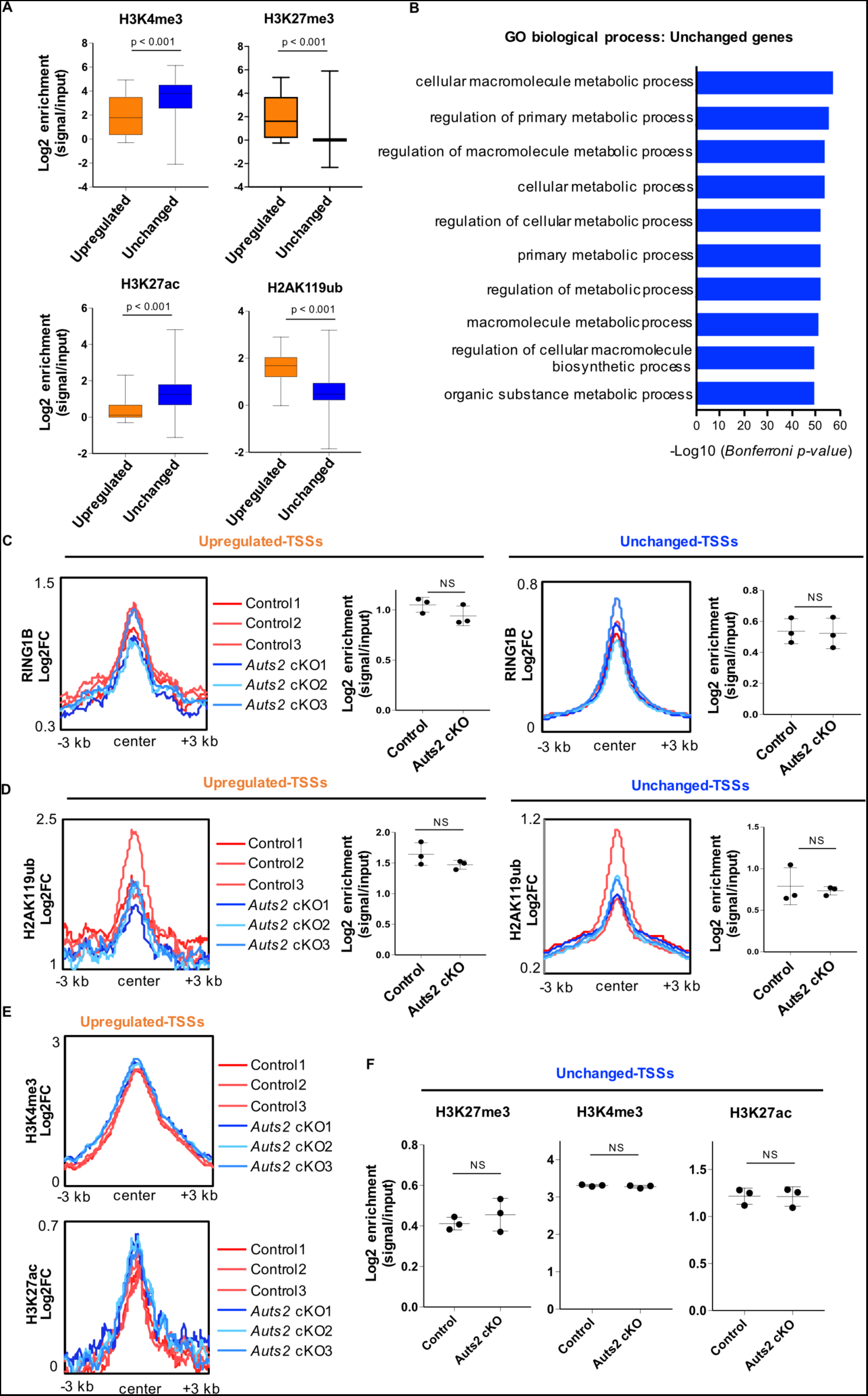
AUTS2 maintains the repressive chromatin status of genes related to neuronal differentiation, as shown in. **Figure 7**. (A) Enrichment of H3K4me3, H3K27me3, H3K27ac and H2AK119ub CUT&Tag signals on IPC-upregulated-TSSs (61 loci) and IPC-unchanged-TSSs (2717 loci). Mann-Whitney U-test. These histone modifications were also significantly different for the other two biological replicates. (B) Graph showing the top 10 GO-term biological processes for genes with IPC-unchanged-TSSs on GREAT. (C) Density plots of RING1B CUT&Tag signals centered on IPC-upregulated-TSSs (left) and IPC-unchanged-TSSs (right) (±3 kb) in control and *Auts2* cKO cells. The graph shows the enrichment of the signals on indicated loci (D) Density plots of H2AK119ub CUT&Tag signals centered on IPC-upregulated-TSSs (left) and IPC-unchanged-TSSs (right) (±3 kb) in control and *Auts2* cKO cells. The graph shows the enrichment of the signals on indicated loci. (E) Density plots of H3K4me3 (top) and H3K27ac (bottom) CUT&Tag signals centered on IPC-upregulated-TSSs (±3 kb) in control and *Auts2* cKO cells. (F) Graphs showing the enrichment of H3K27me3, H3K4me3, and H3K27ac in IPC-unchanged TSSs in control and *Auts2* cKO cells. Data are presented as the mean ± SD (N=three biological replicates). NS, not significant; unpaired Student’s t-test (C, D, F).

**Figure S9:**
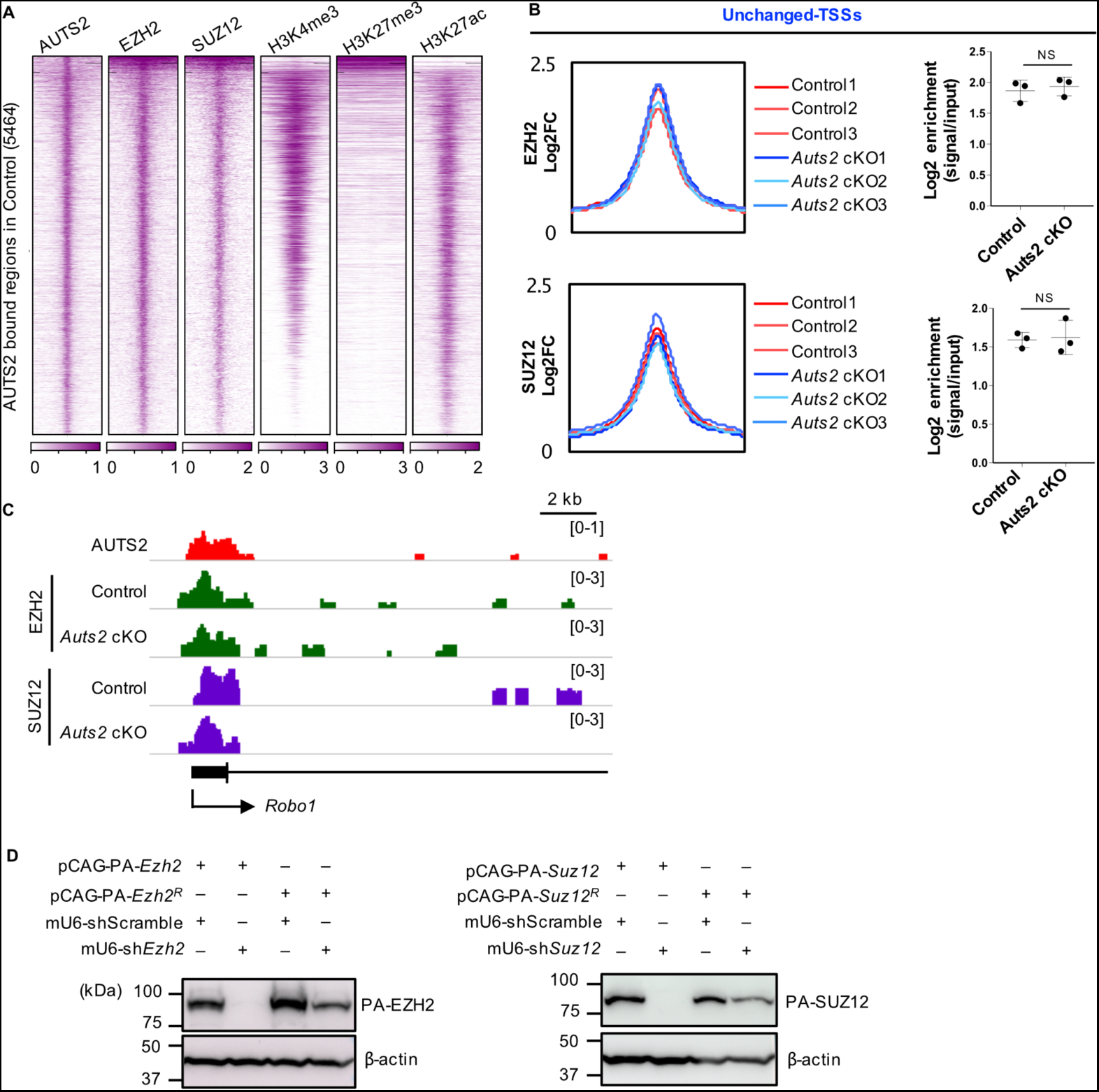
Genome wide distribution of PRC2 proteins in IPCs and KD efficiency to PRC2, as shown in Figure 8. (A) Heatmap of AUTS2, EZH2, SUZ12, H3K4me4, H3K27me3 and H3K27ac CUT&Tag signals centered on AUTS2-binding loci (±3 kb) identified in control cells at E15.5. (B) Density plots of EZH2 (top) and SUZ12 (bottom) signals centered on IPC-upregulated-TSSs (±3 kb) in control and *Auts2* cKO cells. Graphs show the enrichment of EZH2 and SUZ12 on IPC-upregulated-TSSs. Data are presented as the means ± SD (N=three biological replicates). NS, not significant, unpaired Student’s t-test. (C) IGV browser views showing the CUT&Tag signal for AUTS2 in control, EZH2 and SUZ12 in control and *Auts2* cKO cells around the TSS of *Robo1* locus. (D) *In vitro* KD experiments for *Ezh2* (left) and *Suz12* (right) by co-transfection of PA-tagged expression vectors together with scrambled shRNA or indicated shRNA vectors into HEK293T cells. Two days after transfection, the cell lysates were immunoblotted with anti-PA and anti-β-actin antibodies. pCAG-PA-*Ezh2^R^* and pCAG-PA-*Suz12^R^* indicate the shRNA-resistant expression vectors. The KD effect was much lower against each resistant mRNA.

